# Mushroom body output neurons MBONa1/a2 define an odor intensity channel that regulates behavioral odor discrimination learning in larval *Drosophila*

**DOI:** 10.1101/2022.12.14.518530

**Authors:** Abdulkadir Mohamed, Iro Malekou, Timothy Sim, Cahir J. O’Kane, Yousef Maait, Benjamin Scullion, Liria M. Masuda-Nakagawa

## Abstract

The sensitivity of animals to sensory input must be regulated to ensure that signals are detected and also discriminable. However, how circuits regulate the dynamic range of sensitivity to sensory stimuli is not well understood. A given odor is represented in the insect mushroom bodies (MBs) by sparse combinatorial coding by Kenyon cells (KCs), forming an odor quality representation. To address how intensity of sensory stimuli is processed at the level of the MB input region, the calyx, we characterized a set of novel mushroom body output neurons that respond only to high odor concentrations.

We show that a pair of MB calyx output neurons, MBONa1/2, are postsynaptic in the MB calyx, where they receive extensive synaptic inputs from KC dendrites, the inhibitory feedback neuron APL, and octopaminergic sVUM1 neurons, but relatively few inputs from projection neurons. This pattern is broadly consistent in the third instar larva as well as in the first instar connectome. MBONa1/a2 presynaptic terminals innervate a region immediately surrounding the MB medial lobe output region in the ipsilateral and contralateral brain hemispheres. By monitoring calcium activity using jRCamP1b, we find that MBONa1/a2 responses are odor-concentration dependent, responding only to ethyl acetate (EA) concentrations higher than a 200-fold dilution, in contrast to MB neurons which are relatively concentration-invariant and respond to EA dilutions as low as 10^−4^. Optogenetic activation of the calyx-innervating sVUM1 modulatory neurons originating in the SEZ (Subesophageal zone), did not show a detectable effect on MBONa1/a2 odor responses. Optogenetic activation of MBONa1/a2 using CsChrimson impaired odor discrimination learning compared to controls. We propose that MBONa1/a2 form an output channel of the calyx, summing convergent sensory and modulatory input, firing only to high odor concentration, and might affect the activity of downstream MB targets.

## Introduction

To respond adaptively to environmental stimuli, sensory signals defined as sensory objects (Gottfried, 2010), must be encoded in neural representations that are highly selective, to be useful for formation and retrieval of memories. The recognition of sensory objects depends on the selective pattern of activation of a few numbers of neurons at higher centers in the brain, using a combinatorial coding mechanism that is shared across different sensory modalities in insects and mammals (Perez-Orive et al., 2002, Quiroga et al., 2005, Settler and Axel, 2009, DeWeese et al., 2003). However, the dynamic range of intensity of sensory signals can be large. For example in vision, the dynamic range of light intensities is in the order of 10^10^; in olfaction rats can recognize odors over a 50,000-fold range in intensity (Homma et al., 2009), and still a unique image or smell must be recognized across a large range of intensities. The control of gain in the visual pathway is found at many levels, for example by negative feedback of horizontal cells on photoreceptors, and amacrine cells on bipolar cells, the presynaptic cells of ganglion cells. However, how different levels of intensity are encoded and integrated to generate a unique representation of a given sensory object at the higher centers of the brain is not well understood.

The olfactory system shares principles of information processing across insects and mammals, and the numerical simplicity of brains in *Drosophila*, makes it a good model to study olfactory coding. Odors are detected by olfactory sensory neurons (OSNs) in the fly antennae and maxillary palp, or dorsal organ in *Drosophila* larvae. OSNs have different affinities for their ligands, and the combinatorial pattern of their activation is represented in the antennal lobe (AL), the first olfactory center. Here, a nonlinear transformation of intensity coding takes place, weak signals are enhanced and strong signals suppressed (Bhandawat et al., 2007), a principle also found in honeybees (Sachse and Galizia, 2003). Therefore, the AL plays a role in improving signal to noise, adjusting the gain, a process that is regulated by lateral inhibition by the local inhibitory neurons (Asahina et al., 2009). Response tuning of individual OSNs in larvae studied by electrophysiological recordings, showed that individual OSNs respond to different dilutions of an odor, in a range of dilutions over 10^−2^ to 10^−4^, and that the combined responses of OSNs allowed larvae to perceive and discriminate odors in an odor preference test (Kreher et al., 2008). Compared to projection neurons (PNs) that are broadly responsive, Kenyon cells (KCs) in the mushroom body (MB) are selective (Perez-Orive et al., 2002), suggesting a transformation in odor coding between PN input and KC processing. The anatomical organization of KCs in *Drosophila* larval MBs predicts the use of a combinatorial code for odors, generating an odor identity channel (Masuda-Nakagawa et al., 2005). KCs are quiescent and respond with one or 2 spikes to PN input, and their responses are relatively concentration-invariant (Stopfer et al., 2003, Ito et al., 2008). In the mammalian piriform cortex, odor identity is suggested to be encoded in a subset of odor concentration-invariant piriform cortex neurons, while piriform cortex neurons can respond to a 100-fold concentration range with different odor representations (Roland et al., 2017).

To understand the neural circuitry that regulates the selectivity of sensory representations, and its regulation by odor intensity, we use the *Drosophila* larval MB calyx. MBs are centers for associative learning in the insect brain, and the calyx is the dendritic input region with a role in odor discrimination. The calyx receives stereotypic input from projection neurons (PNs) and is innervated by a feedback inhibitory neuron, the larval APL (Masuda-Nakagawa et al., 2005, 2014), and two octopaminergic neurons called sVUM1 originating in the SEZ, which can modulate behavioral odor discrimination (Wong, Wan et al., 2021). Two output neurons, named MBONa-1 and MBONa-2, arborize in the MB calyx (Saumweber et al., 2018).

Odd neurons are a group of eight neurons (Slater et al., 2015) that include two neurons named MBONa1/a2 (Saumweber et al., 2018) that widely innervate the calyx of the MBs, and have been proposed to receive multiple PN inputs (Slater et al., 2015). Odors can be discriminated by a combinatorial mechanism at the MB calyx, that allows the representation of many odors (Masuda-Nakagawa et al., 2005). The balance between discrimination and sensitivity affects learning; higher sensitivity could improve learning, while lower sensitivity could decrease learning, while improving discrimination. Slater et al., (2015) showed that Odd neurons enhanced the ability to discriminate different concentrations of odors in a chemotaxis assay; however, learning was not tested.

We have shown previously that the calyx-innervating Odd neurons, MBONa1/a2, labeled by *GAL4-OK263*, have potential synaptic contacts with both octopaminergic sVUM1 neurons, sVUMmd1 and sVUMmx1 (Wong, Wan et al., 2021), and therefore MBONa1/a2 might be subject to modulation by OA input.

Here we characterize the connections and polarity of MBONa1/a2 neurons in the larval calyx. We show that MBONa1/a2 respond to odor in a concentration-dependent manner, in contrast to the relatively concentration-invariant KCs. They also affect learning performance, although not in a strongly concentration-dependent manner. MBONa1/a2 may nonetheless represent a channel that conveys odor intensity information to the output region of the MBs, and therefore could potentially have a role in concentration-dependent modulation of MB output neurons and signals.

## Materials and Methods

### Fly stocks

All stocks were maintained on cornmeal-yeast-agar medium at 25**°**C in a 12-hour day/night cycle. Stocks are listed in Table 1.

**TABLE 1.**
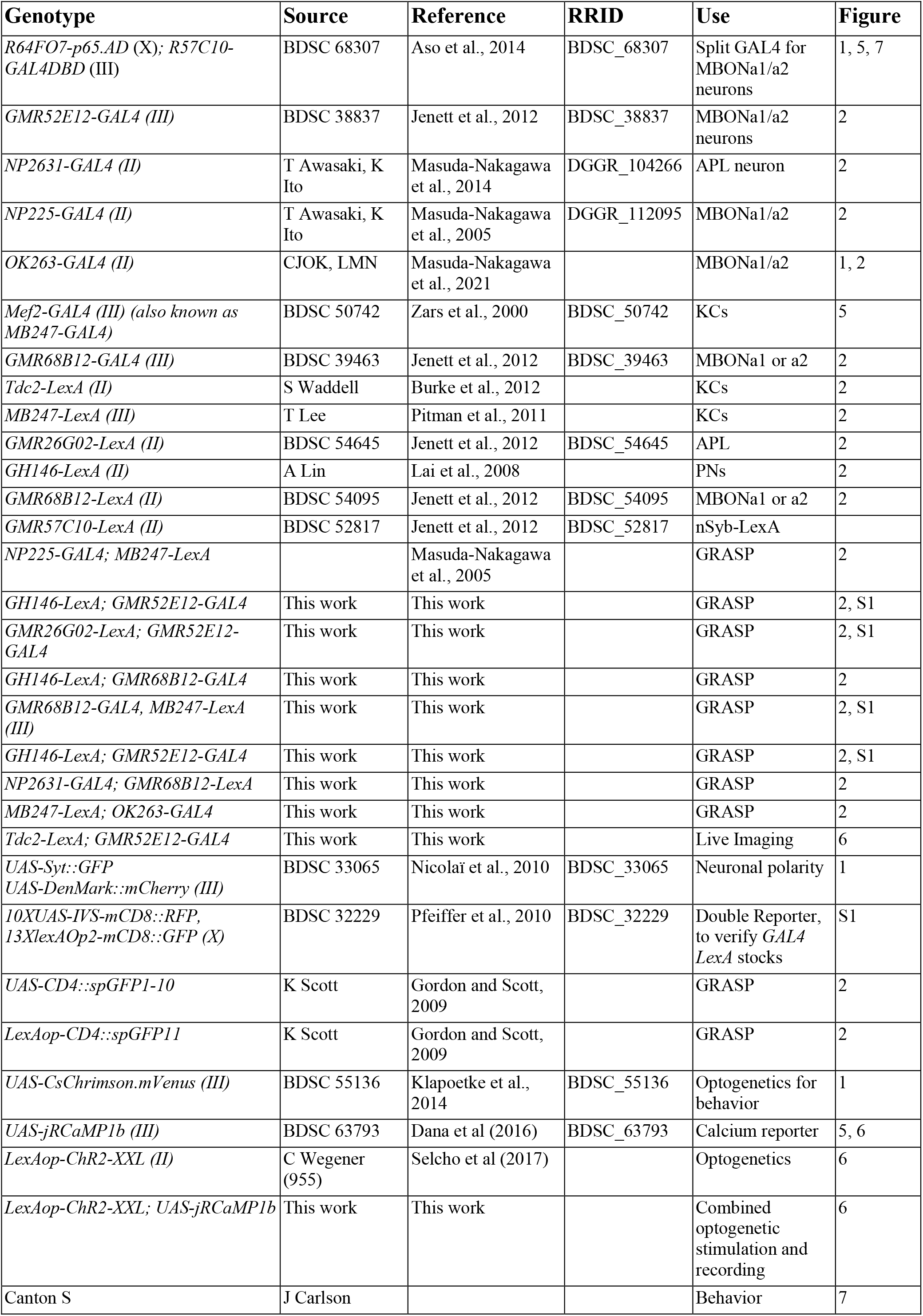
*Drosophila* stocks used. Includes all genotypes used for this study, including those that do not appear in figures.

| Genotype | Source | Reference | RRID | Use | Figure |
| --- | --- | --- | --- | --- | --- |
| *R64FO7-p65.AD (X); R57C10-GAL4DBD (III)* | BDSC 68307 | Aso et al., 2014 | BDSC_68307 | Split GAL4 for MBONa1/a2 neurons | 1, 5, 7 |
| *GMR52E12-GAL4 (III)* | BDSC 38837 | Jenett et al., 2012 | BDSC_38837 | MBONa1/a2 neurons | 2 |
| *NP2631-GAL4 (II)* | T Awasaki, K Ito | Masuda-Nakagawa et al., 2014 | DGGR_104266 | APL neuron | 2 |
| *NP225-GAL4 (II)* | T Awasaki, K Ito | Masuda-Nakagawa et al., 2005 | DGGR_112095 | MBONa1/a2 | 2 |
| *OK263-GAL4 (II)* | CJOK, LMN | Masuda-Nakagawa et al., 2021 |  | MBONa1/a2 | 1, 2 |
| *Mef2-GAL4 (III) (also known as MB247-GAL4)* | BDSC 50742 | Zars et al., 2000 | BDSC_50742 | KCs | 5 |
| *GMR68B12-GAL4 (III)* | BDSC 39463 | Jenett et al., 2012 | BDSC_39463 | MBONa1 or a2 | 2 |
| *Tdc2-LexA (II)* | S Waddell | Burke et al., 2012 |  | KCs | 2 |
| *MB247-LexA (III)* | T Lee | Pitman et al., 2011 |  | KCs | 2 |
| *GMR26G02-LexA (II)* | BDSC 54645 | Jenett et al., 2012 | BDSC_54645 | APL | 2 |
| *GH146-LexA (II)* | A Lin | Lai et al., 2008 |  | PNs | 2 |
| *GMR68B12-LexA (II)* | BDSC 54095 | Jenett et al., 2012 | BDSC_54095 | MBONa1 or a2 | 2 |
| *GMR57C10-LexA (II)* | BDSC 52817 | Jenett et al., 2012 | BDSC_52817 | nSyb-LexA |  |
| *NP225-GAL4; MB247-LexA* |  | Masuda-Nakagawa et al., 2005 |  | GRASP | 2 |
| *GH146-LexA; GMR52E12-GAL4* | This work | This work |  | GRASP | 2, S1 |
| *GMR26G02-LexA; GMR52E12-GAL4* | This work | This work |  | GRASP | 2, S1 |
| *GH146-LexA; GMR68B12-GAL4* | This work | This work |  | GRASP | 2 |
| *GMR68B12-GAL4, MB247-LexA (III)* | This work | This work |  | GRASP | 2, S1 |
| *GH146-LexA; GMR52E12-GAL4* | This work | This work |  | GRASP | 2, S1 |
| *NP2631-GAL4; GMR68B12-LexA* | This work | This work |  | GRASP | 2 |
| *MB247-LexA; OK263-GAL4* | This work | This work |  | GRASP | 2 |
| *Tdc2-LexA; GMR52E12-GAL4* | This work | This work |  | Live Imaging | 6 |
| *UAS-Syt::GFP*<br>*UAS-DenMark::mCherry (III)* | BDSC 33065 | Nicolai et al., 2010 | BDSC_33065 | Neuronal polarity | 1 |
| *10XUAS-IVS-mCD8::RFP, 13XlexAOp2-mCD8::GFP (X)* | BDSC 32229 | Pfeiffer et al., 2010 | BDSC_32229 | Double Reporter, to verify *GAL4 LexA* stocks | S1 |
| *UAS-CD4::spGFP1-10* | K Scott | Gordon and Scott, 2009 |  | GRASP | 2 |
| *LexAop-CD4::spGFP11* | K Scott | Gordon and Scott, 2009 |  | GRASP | 2 |
| *UAS-CsChrimson.mVenus (III)* | BDSC 55136 | Klapoetke et al., 2014 | BDSC_55136 | Optogenetics for behavior | 1 |
| *UAS-jRCaMP1b (III)* | BDSC 63793 | Dana et al (2016) | BDSC_63793 | Calcium reporter | 5, 6 |
| *LexAop-ChR2-XXL (II)* | C Wegener (955) | Selcho et al (2017) |  | Optogenetics | 6 |
| *LexAop-ChR2-XXL; UAS-jRCaMP1b* | This work | This work |  | Combined optogenetic stimulation and recording | 6 |
| Canton S | J Carlson |  |  | Behavior | 7 |

### Immunohistochemistry and confocal imaging

Third-instar wandering larvae (144-176 hours AEL) were dissected in cold PBS, fixed in 4% Formaldehyde / PEM buffer (0.1 M PIPES; 2 mM EGTA; 1 mM MgSO_4_; NaOH), PH 7.3, for 2 hours at 4°C, washed for 3×10 minutes (or 4×15 minutes) in 0.3% Triton-X in PBS (PBT) and incubated in 10% NGS (Normal goat serum) in 0.3% PBT for 1 hour at room temperature. Brains were incubated in primary antibody in 10% NGS-0.3% PBT at 4°C for 2-3 days on a mini disk rotor (Biocraft, BC-710), washed for 3×15 minutes with 0.3% PBT and further incubated in secondary antibody in 10% NGS at 4°C for 2-3 days again on the mini disk rotor. Brains were finally washed 1×15 minutes with PBT, followed by 3×10 minutes with PBS, and left in 50% Glycerol/PBS at 4°C for at least one night prior to imaging. Primary and secondary antibodies are listed in Table 2, other reagents in Table 3. Imaging was carried out using an SP8 Confocal Microscope with a 40X NA1.3 water objective.

**TABLE 2.**
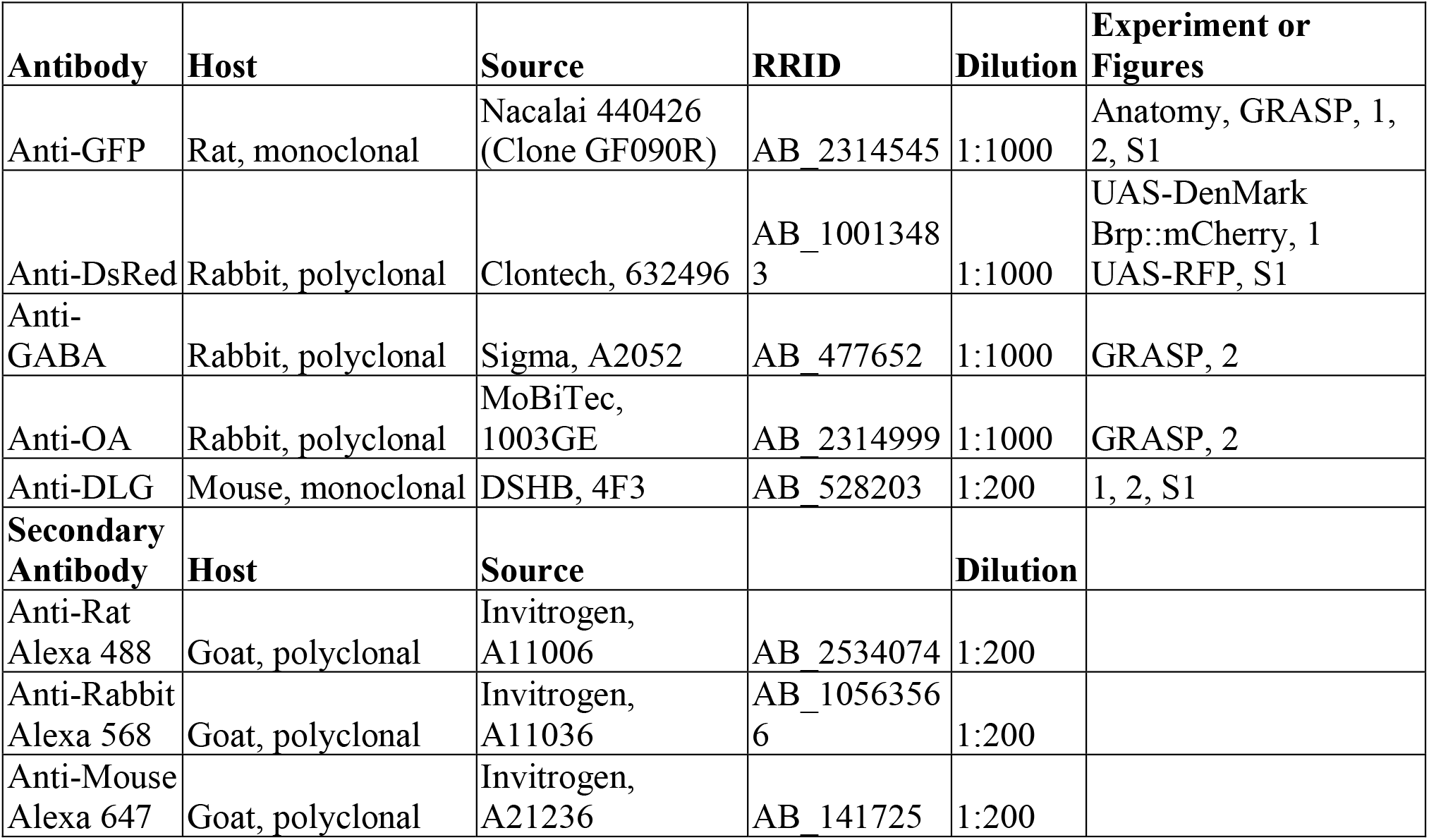
Antibodies.

**TABLE 3.** Other reagents.

| **Reagent** | **Source** | **Experiment or Figures** |
| --- | --- | --- |
| PBS | Sigma P4417 | Anatomy, GRASP |
| Formaldehyde | Polysciences 18814 | Anatomy, GRASP |
| PIPES | Sigma P1851 | Anatomy, GRASP |
| EGTA | Sigma E3889 | Anatomy, GRASP |
| Triton-X | Sigma T8787 | Anatomy, GRASP |
| Normal goat serum | Vector Labs S-1000 | Anatomy, GRASP |
| Paraffin oil | Sigma-Aldrich, 76235 | Live imaging, Behavior |
| Pentyl Acetate | Sigma-Aldrich, 109584 | Live imaging, Behavior |
| Ethyl Acetate | Sigma-Aldrich, 319902 | Live imaging, Behavior |
| Ethanol | Sigma-Aldrich, 32221 | Behavior |
| all-trans-retinal (ATR) | Sigma-Aldrich, R2500 | Live imaging, Behavior |
| D-(-)-Fructose | Sigma-Aldrich, 47740 | Behavior |
| Agarose | Sigma-Aldrich, A9539 | Behavior |

### CATMAID

The publicly available first-instar larval connectome on the CATMAID software on the Virtual Fly Brain site (https://l1em.catmaid.virtualflybrain.org; Licence CC-BY-SA_4.0, Saalfeld et al., 2009, Schneider-Mizell et al., 2016) was used to study neuron morphology and synaptic partners. The ‘Connectivity Widget’ was used to find the listed presynaptic partners of MBONa1/a2. For each presynaptic partner, the location of each of the synapses made with MBONa1/a2 was noted by clicking on the individual synapse connector number and looking at the location shown on the 3D reconstruction of the neuron on the “3D viewer”. The number of synapses made by each type of neuron with MBONa1/a2 neuron in the calyx were counted and graphs were generated using Microsoft Excel version 16.60. For each of the MBONa1/a2 neurons, the number of cyan (postsynaptic) and red (presynaptic) sites was counted three times and an average was calculated (Supplementary Table 1).

### Live imaging and optogenetics

Live imaging was performed as previously described (Masuda-Nakagawa et al., 2009). Wandering stage L3 *Drosophila* were dissected and mounted for imaging under an Olympus BX50-WI microscope with a Zeiss W Plan-Apochromat 40x/1.0 DIC M27 objective and using an Andor iXon + DU-88E-CO-#BV EM-CCD camera (Andor, Belfast, UK), via a Cairn Research Optosplit II (Faversham, UK).

Wandering stage L3 larvae for combined optogenetics and imaging were dissected under dim amber (591 nm) light and the condenser light of the BX50-WI was passed through an ET632/60 M emission filter. A Cairn OptoLED LED mount (Cairn Research) on the Olympus BX50-WI BX-FLA vertical illuminator was used to illuminate the sample through the objective. The vertical illuminator aperture was minimized to a diameter of approximately twice that of the calyx (from a dorsal view) and centered within the camera’s field of view. A 470nm LED and power supply (Cairn Research, OPTOLED Light source) with an LED mount (Cairn Research, LED mount) was used alongside Polygon400 (polygon400-G) for patterned illumination. Polyscan2 software (Mightex) was used to control the illumination pattern.

Throughout the optogenetics experiment, the full camera capture region was illuminated (256×256 µm). A microscope slide power sensor (Thorlabs S170C) with an optical power and energy meter (Thorlabs PM400) was used to calibrate the LED and measure the LED power at the specimen, *Tdc2-lexA* was crossed to *lexAop-jRCamP1b, lexAop-ChR2(XXL)*, and it was subsequently shown that 470nm LED of 8.55 µW intensity applied for 500ms was the optimal stimulus strength and duration to evoke a strong increase in calcium with minimum light exposure. A T495lpxr dichroic mirror (Chroma, VT, USA) directed the LED light onto the sample while preventing 470nm light reflected off the sample from reaching the microscope output.

The shutter of a Yokogawa CSU22 spinning disc confocal (Yokogawa Electric Corporation) was controlled by Micro Manager (Edelstein et al., 2010), and the PC interface with the spinning disc was via the multifunction I/O card (National Instruments, PCI-6221). The camera interface was via an Andor interface card (Andor, PCI controller) and Metamorph software (Metamorph meta imaging series version 7.0) was used to control the camera settings and image acquisition. The Cairn Research (Faversham, UK) laser controller was used to deliver a 561nm excitation laser, at 20% laser power; at this power, *LexAop-ChR2-XXL/nSyb-LexA; UAS-jRCaMP1b/ MKRS* larvae showed no movement in response to the laser. The room was kept at 23°C. A Master-8-cp controller (AMPI, Jerusalem, Israel) synchronized the timing of imaging, LED illumination, and odor delivery.

Humidified odors were presented through valves controlled by the Master-8-cp controller as described (Masuda-Nakagawa et al., 2009). All crosses were performed on cornmeal-yeast-agar medium, and for optogenetics media were supplemented with 100 μM all-trans-Retinal (Sigma, R2500). Crosses were kept at 23°C in the dark wrapped in tinfoil, and when necessary handled under dim amber light (591 nm).

Combined optogenetics and activity imaging was performed by crossing *LexAop-ChR2-XXL; UAS-jRCaMP1b/TM6B* virgin females to a stock carrying *52E12-GAL4* and *Tdc2-LexA* insertions, and collecting wandering third instar progeny. ChR2-XXL function was confirmed by crossing female parents to *nSyb-LexA/CyO::GFP* males and testing undissected, non-*CyO::GFP* larval progeny for light-induced body contraction under imaging conditions. Driver genotypes in all *GAL4* and *LexA* combination lines were tested by imaging RFP and GFP expression in the larval progeny of a cross between males of each line to virgin females of a *UAS-RFP LexAop-GFP* double reporter line (Bloomington stock 32229).

### Paradigms

Concentration-dependence of MBONa1/a2 and KC responses: a pulse of EA at different concentrations diluted in mineral oil was applied for 2 seconds, at 2 seconds after the start of image acquisition. Image acquisition was at a frame rate of 5 frames/second, for 100 ms. Image acquisition continued for a total of 12 seconds. Thirty-second intervals with no laser exposure were introduced between image acquisitions to re-establish baseline fluorescence. The average fluorescence of the first 10 frames of image acquisition was taken as baseline.

Combined optogenetics and imaging experiment: “odor only” consisted of 3 seconds of baseline image acquisition, followed by a 10-fold diluted EA pulse for 2 seconds. Frame rate and laser duration were the same as above. Image acquisition was for a total of 13 seconds. The average fluorescence value of the first 10 frames was taken as baseline. “Odor+Light” consisted of 2 seconds of baseline image acquisition, followed by a pulse of 470nm LED for 500 ms, then an odor pulse of a 10-fold dilution of EA was applied at 3 seconds after the start of image acquisition for 2 seconds, with a total image acquisition time of 13 seconds. An interval of 76 seconds between the end of one acquisition and the start of the next was introduced to allow time for baseline recovery, according to τ_off_ for ChR2-XXL being 76 seconds (Dawydow et al., 2014). Frame rate and laser duration were the same as above. Pulse duration was determined by titrating time and recording increase in fluorescence in *Tdc2* neurons expressing *ChR2-XXL* and *jRCAMP1b*, near the primary processes where the signal was stronger. “Light only” was as above except that no odor pulse was applied. In control experiments using *52E12-GAL4* and *UAS-JRCaMP1b*, in the presence of a *LexAop-ChR2-XXL* construct without the *Tdc2* driver, five pulses of 470nm LED of 50 ms were applied at 200-ms intervals for a total time of 1 second for one of two sets of experiments instead of a single 500-ms light pulse as above, but no significant differences between the two light treatments were detected in the peak responses obtained (Mann-Whitney test or t test as appropriate), and therefore these data were merged.

### Image analysis

ImageJ (Schindelin et al., 2012) was used to analyze the stacks generated by Metamorph, and to obtain the standard deviation of all pixels in the recording area. A region of interest (ROI) was selected to correspond to the responding pixels with highest standard deviation, ensuring the same ROI was obtained when comparisons were to be made between different stimulation conditions. The Z-axis profile function was then used to quantify the fluorescence changes over time which were subsequently saved as Excel files. The same ROI outline was then moved to an adjacent, non-responding region and the procedure was repeated to calculate background fluorescence over time. After subtracting background fluorescence from each frame, baseline fluorescence calculated as the average of the first 10 frames before odor delivery, were subtracted from each frame value to obtain ΔF. Normalized signal intensity ΔF/F at each time point was calculated by dividing ΔF over baseline fluorescence (Masuda-Nakagawa et al., 2014). To ensure these changes were not due to the order in which concentrations were tested (possibility for adaptation affecting the results for the later tested concentrations), randomization of the order in which the concentrations were tested was also practiced. In this experiment and subsequent experiments, data from larvae that had moved during imaging were excluded as it was impossible to tell if the fluorescence change was due to movement or an odor response. Each acquisition consisted of 60 frames of background-subtracted intensity data acquired over 12 seconds. To smoothen out frame-to-frame variation, we used the overlapping moving averages of three sequential frames, leaving 58 frames of smoothed data.

### Statistics

Analysis was carried out using GraphPad Prism 8 software, and statistical tests used for each experiment are shown in figure legends. Statistical significance was defined as P<0.05 throughout, taking account of post-hoc and multiple testing.

### Behavioral assay

Larval culture. For testing behavioral roles of MBONa1/a2, flies of genotype *w; 20xUAS-IVS-CsChrimson.mVenus (III)* were crossed to either *w, R64F07-p65-AD* ; *R57C10-GAL4-DBD* for MBONa1/a2 behavior assay, or Canton S as positive control. Larvae were allowed to develop in food vials containing 100 μM all-trans-retinal, in the dark at 23 degrees. Adults were transferred into new vials both in the morning and in the evening, and progeny collected after 136-152 hours at 23°C.

Behavioral arena. Agarose petri dishes were prepared the day before use, by using 100 ml of distilled water with 0.9% agarose (Sigma A9539). Fructose petri dishes were prepared similarly, but containing 35% fructose (Sigma-47740). Petri dishes had perforated lids, to facilitate creation of odorant gradients within the dish, by sucking air from a benchtop fume extractor (Sentry Air Systems (SAS), SS-200-WSL) at the back of the assay platform. Odorants were diluted in paraffin oil (Sigma-Aldrich 76235) and 10µl was added to custom-built Teflon containers with pierced lids (7 holes), on the surface of agarose plates.

Light apparatus. The light apparatus contained a BK Precision DC power pack, connected to a pulse generator, driving four sets of amber light LEDs (591 nm), Luxeon star Amber LED on Tri-Star Base, 330 lm at 350 mA (SP-03-A5). The irradiance on the platform was 0.06 μW/mm2 on average (24 μW on a 20 × 20-mm sensor) on the 8.5-cm plate. the pulse generator was constructed as described by deVries and Clandinin (2013), by the Psychology Department workshop of the University of Cambridge. One cycle of pulses consisted of 10-ms pulses at 10Hz for 30 s, followed by 30 s without pulses. This cycle was repeated 5 times, making a conditioning step of 5 minutes.

Behavior conditioning. Third-instar larvae were collected from vials using a metal sieve, and washed with tap water. Larvae were washed twice by transferring through a drop of water, and then placed on the conditioning agarose plate (35% fructose). A Teflon container with 10 μl of odor A to be reinforced was placed on the left side, and another containing paraffin oil (neutral) symmetrically on the right side, at positions marked on a black background, used to facilitate visualization of the larvae. Larvae were conditioned on the fructose plate with odor A for 5 minutes under weak blue light. Larvae were then transferred to a water droplet using a brush, and again to a second water droplet to ensure no fructose traces remained, and then to an agarose plate lacking fructose, on which a Teflon container with 10 μl of odor B (non-reinforced) was placed on the left side and a container with paraffin oil (neutral) on the right side. Larvae were conditioned for 5 minutes under weak blue light as above. This conditioning procedure on both plates was repeated for three cycles. For experiments using activation of OA neurons, the entire conditioning cycles were carried out under amber light.

Odor dilutions. Dilutions of ethyl acetate (EA) at 1:2000 and pentyl acetate (PA) at 1:500, and EA at 1:4000 and PA at 1:1000 diluted in mineral oil were used.

Testing. Larvae were tested by placing them on an agarose plate carrying a container with EA on one side, and a container with PA on the other. Test was under blue light for 5 minutes. Larvae were counted on the side of the conditioned odor, the unconditioned odor or in the neutral zone in the middle, and the performance index (PI) was calculated, using the formula:

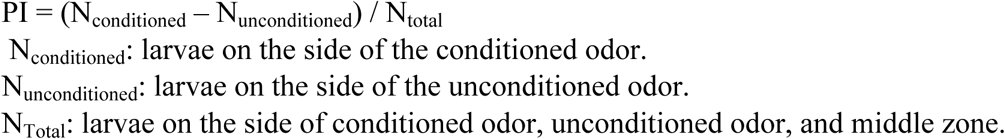

Learning Index (LI) was calculated after both a conditioning with odor A and a reciprocal conditioning with odor B (with a different sample of larvae), using the formula:

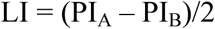

Statistical analysis. A two-way ANOVA using Graph Pad Prism software was used to test whether learning index was affected by odor concentration or light.

## Results

### MBONa1/a2 are postsynaptic in the MB calyx

Odd neurons defined as a group of 8 neurons, labeled by *Odd-GAL*, some of which innervate the larval calyx (Slater et al., 2015). Connectomic analyses of the first instar larva showed that two of them innervate the calyx, and these were named as MBON-a1 and MBON-a2 (Saumweber et al., 2018). In the 3^rd^ instar larva, two neurons with cell bodies located posteriorly to the MBs are labeled in the split-GAL4 line *R64FO7-p65.AD* (X); *R57C10-GAL4DBD* (III) driving expression of *UAS-CsChrimson.mVenus* (Fig. 1A). They join in a primary process that bifurcates anteriorly to the calyx, with one branch penetrating the calyx ventrally and branching into an extensive network throughout the calyx, and the main process extending anteriorly to innervate a region wrapping around the MB medial lobes (ML). One axon leaves the area surrounding the ML to cross the midline and innervates the equivalent regions around the contralateral ML (Fig. 1A, arrow). The innervation of the calyx is dense but localized to interglomerular spaces and the core of the calyx, avoiding the calyx glomeruli, labeled by anti-DLG, that are neuropiles with connectivity between PN boutons and KC dendritic claws Fig 1B, Supplementary Fig. 1A, B. However, MBONa1/a2 show extensive overlap with the GABAergic larval APL in the calyx (Supplementary Fig. 1C). Processes in the calyx show thin processes as well as thick enlargements. GABA is in the vicinity of MBONa1/a2 processes, but not in the vicinity of ACh-labeled PN boutons (data not shown). In the MB medial lobe, axons form a circle around it but do not innervate the lobes (Fig. 1C). MBONa1/a2 innervation of the calyx is mainly postsynaptic as shown by expression of the dendritic marker DenMark::mCherry (Nikolai et al., 2010), driven by *OK263-GAL4* (Wong, Wan et al. 2021), only a few puncta were seen using Syt::GFP; however, strong Syt::GFP labeling but not DenMark was observed around the region of the ML (Fig. 1D), implying that these MBONa1/a2 projections are presynaptic. Consistent with this, we also found strong localization of a second presynaptic marker, nSyb::GFP at the output regions around the ML (Supplementary Fig. 1D).

**FIGURE 1.**
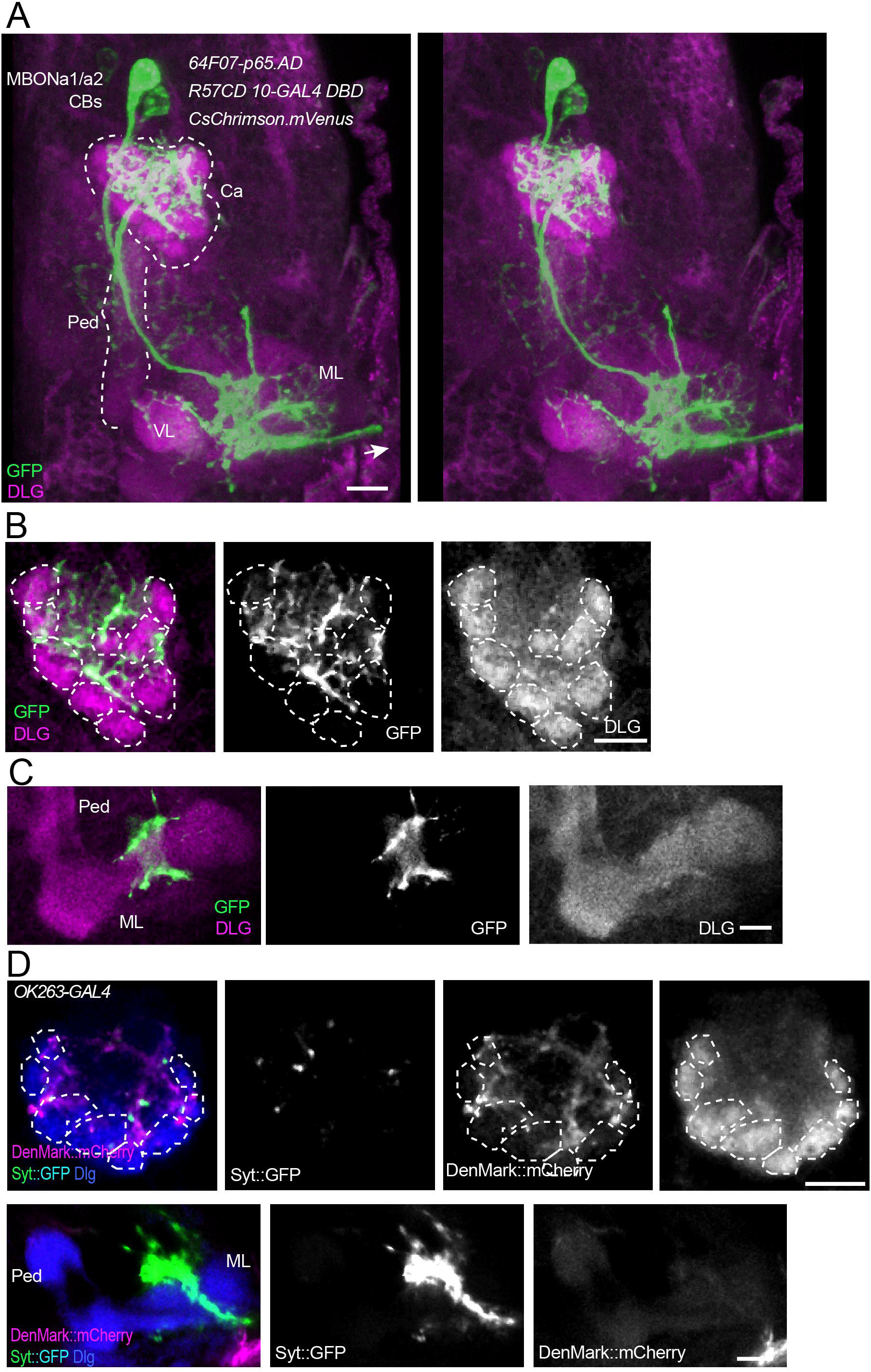
MBONa1/a2 neurons innervate the calyx and output regions around the MB medial lobes. (**A)** A 3D stereo pair of a dorsal view of the MB in the right brain hemisphere showing a pair of MBONa1/a2 neurons labeled by the split-GAL4 line *64FO7-p65.AD* (X); *R57C10-GAL4DBD* (III) driving *UAS-CsChrimson.mVenus*. mVenus is labeled by anti-GFP, and MBs are labeled by anti-Dlg. Notice the extensive innervation of the calyx, and of the region surrounding the vertical lobe (VL), and medial lobe (ML) of the MBs. Both MBONa1/a2 neurons also project to the contralateral brain hemisphere (arrow). (**B)** A single section of the calyx shown in A, showing the dense innervation by MBONa1/a2 throughout the calyx core and interglomerular spaces. (**C)** A single confocal section showing MBONa1/a2 innervation of a region in the vicinity of the medial lobe of the MBs, from a ventral section of the preparation shown in A. (**D)** Projections of confocal sections of MBONa1/a2 expressing DenMark::mCherry and Syt::GFP under the control of *OK263-GAL4*, detected by antibodies to DsRed and GFP respectively. Top row, calyx; bottom row, ML and pedunculus. Anterior is to the bottom in all panels. Glomeruli indicated by dotted lines. Ca, calyx; Ped, pedunculus; VL, vertical lobe; ML, medial lobe. Scale bars, 10 µm.

### Pattern of calyx innervation by calyx-innervating neurons

Third instar larvae of the genotypes used for GRASP analysis were used to drive the double reporter *UAS-mCD8::RFP LexAop-mCD8::GFP*, to visualize the pattern of calyx innervation. *MB247-LexA*, which drives expression in KCs, shows dense labeling of the calyx, whereas MBONa1/a2 expressing *OK263-GAL4* innervate the core of the calyx and interglomerular space, Supplementary Fig. 1A. The dense innervation of the calyx by KCs is compatible with the diffuse labeling observed by GRASP (Fig. 2A, Bi).

**FIGURE 2.**
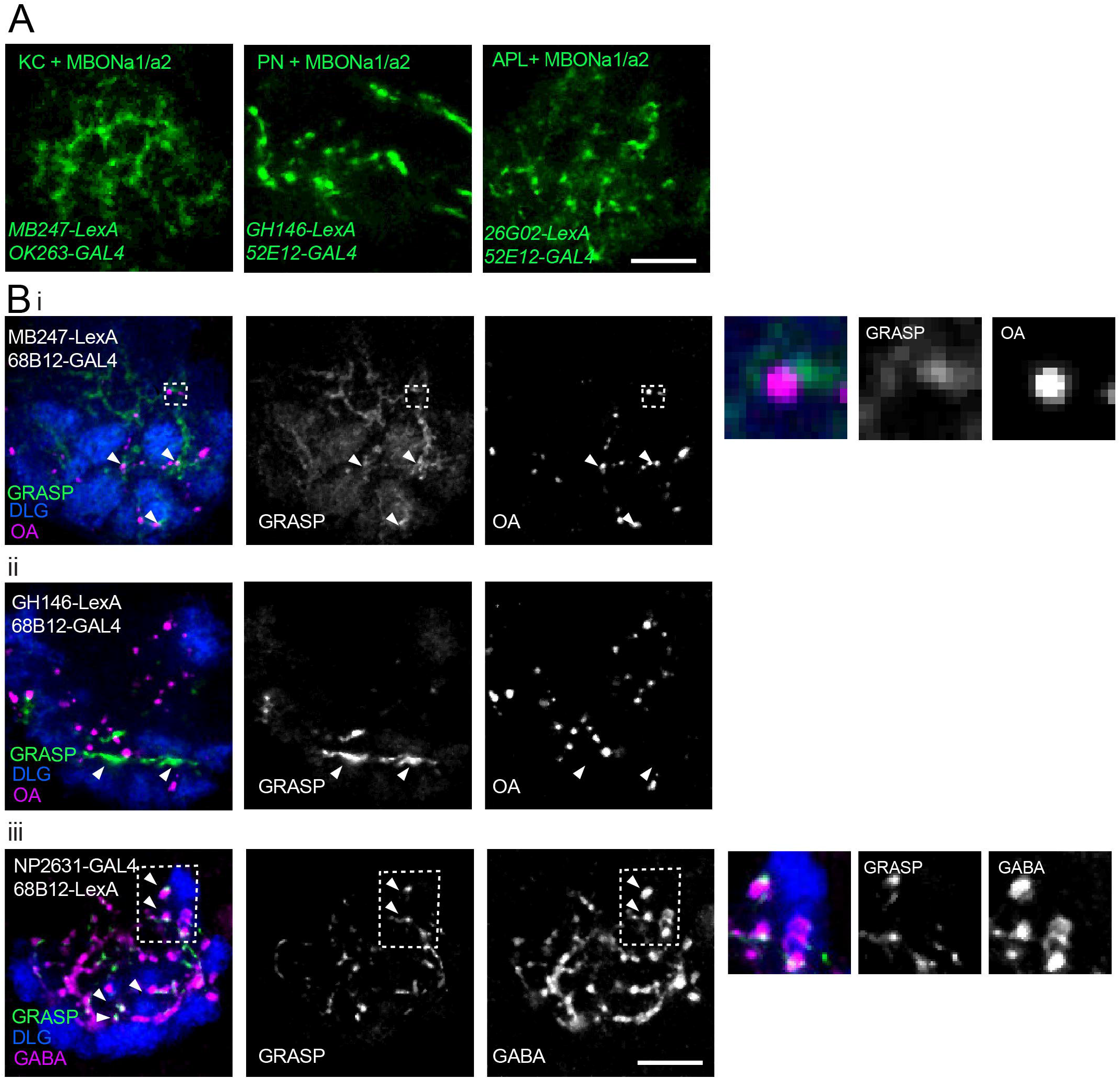
Connectivity of MBONa1/a2 and other calyx neurons in third instar larvae. (**A)** Live GRASP signals of MBONa1/a2 with dendrites of KCs, axonal termini from PNs, or axonal processes of larval APL. A line carrying GRASP constructs *UAS-CD4::spGFP1-10* and *LexAop-CD4::spGFP11*, was crossed to flies containing the *GAL4* and *LexA* constructs shown. GRASP signals were detected in the 3^rd^ instar wandering stage larval progeny. GRASP signal was detected as native GFP fluorescence in z-projections of a few confocal sections. All panels are right brain images, anterior to bottom. Scale bar 10µm. **(B)** GRASP signals between one of the MBONa1/a2 neurons and KCs, PNs, or APL visualized by immunolabeling. A line carrying GRASP constructs *UAS-CD4::spGFP1-10* and *LexAop-CD4::spGFP11* was crossed to flies containing the constructs shown. GRASP signals were detected in the 3^rd^ instar wandering stage larval progeny using monoclonal rat anti-GFP. **(i)** In MBONa1/a2-KC GRASP, arrowheads indicate GRASP signal in proximity to OA boutons. Inset inside broken line is shown enlarged to the right of the panels. **(ii)** In MBONa1/a2-PN GRASP, GRASP signal is observed in PN axonal tracts and may not represent synaptic contacts (arrowheads). **(iii)** MBONa1/a2-APL: arrowheads indicate co-localization of GABA and GRASP signals. Inset inside broken line is shown enlarged to the right of the panels. Calyx glomeruli are visualized using anti-Dlg. Right brain calyces are shown, anterior is to the bottom. Scale bar: 10 µm.

Axonal tracts of PNs can be observed traversing the calyx and ending in boutons, while a dense labeling of process within the calyx core avoiding glomerular regions is observed in larvae carrying *GH146-LexA* and *52E12-GAL4*. This suggests that the GRASP GFP signal observed is likely to be due to proximity of axons from PNs and MBONa1/a2 dendrites (Supplementary Fig. 1B), and that MBONa1/a2 may not receive direct synaptic inputs from PNs.

The pattern of innervation of MBONa1/a2 and the larval APL expressing *UAS-mCD8::RFP LexAop-mCD8::GFP*, in larvae carrying *26G02-LexA* and *52E12-GAL4*, shows extensive overlap of processes, and contacts around APL boutons (Supplementary Fig. 1C).

### Extensive contacts of MBONa1/a2 with calyx neurons

To analyze the connectivity between MBONa1/a2 and other neurons in the calyx, we used *GAL4* lines expressing in MBONa1/a2, and *LexA* lines expressing in potential partner neurons in the calyx, to express GRASP (GFP Reconstitution Across Synaptic Partners, Gordon and Scott, 2008) constructs *UAS-CD4::spGFP1-10* and *LexAop-CD4::spGFP1* in third instar larval brains. We used a number of *GAL4* lines to label MBONa1/a2 (Fig. 2. Supplementary Fig. 1) mainly due to the need for different chromosomal locations for stock construction. Extensive native GRASP GFP fluorescence was observed between MBONa1/a2 on the one hand, and KCs, PNs, and APL on the other (Fig. 2A). KC-MBONa1/a2 GRASP showed diffuse labeling throughout the calyx with some puncta, MBONa1/a2-PN GRASP showed strong labeling along axonal tracts, and MBONa1/a2-APL GRASP showed characteristic GFP puncta.

We also confirmed the occurrence and localization of GRASP signals in the 3^rd^ instar wandering stage larvae by antibody labeling. Brains were immunolabeled to visualize glomeruli by anti-DLG (Lahey et al., 1994), GFP by rat monoclonal anti-GFP, and subsets of synaptic sites were visualized by anti-octopamine (OA) for MBONa1/a2-KC or MBONa1/a2-PN connectivity, and anti-GABA for MBONa1/a2-APL connectivity. KC-MBONa1/a2 GRASP showed a diffuse pattern with some GRASP signals localized in the vicinity of OA boutons, suggesting a synaptic localization for some of the signals, consistent with the MBONa1/a2 contacts seen previously using GRASP (Wong, Wan et al., 2021). These were in the calyx core, or interglomerular space (Fig. 2Bi, arrows and inset). MBONa1/a2-PN GRASP showed a strong signal that we interpret as axonal tracts because of their length along the borders with glomeruli. It is hard to determine whether some synaptic contacts were present between PN axonal tracts and MBONa1/a2 dendrites (Fig. 2B ii). MBONa1/a2-APL GRASP showed the characteristic GRASP puncta, and their synaptic localization is shown by co-localization with GABA. Many GRASP signals are also observed in the core of the calyx and in the interglomerular space (Fig. 2Biii).

### Pattern of innervation of MBONa1/a2 in the first instar larva

In the connectome of the first instar larva, a pair of calyx-innervating mushroom body output neurons have been annotated in the CATMAID tool as MBON-a1-R/ MBE7a right, and MBON-a2-R/ MBE7b in the larval right brain hemisphere, and another pair MBON-a1-L/ MBE7a left and MBON-a2-L/ MBE7b in the left brain hemisphere (Supplementary Table 1 from Eichler et al. 2017). Processes of MBON-a1-R/a2-R (Fig. 3Ai) project to the calyx, where processes are predominantly postsynaptic (Fig. 3Bi-iii), and then to the output region in the ipsilateral and contralateral regions around the MB medial lobes containing both postsynaptic as well as presynaptic sites (Fig. 3C). There was extensive regional overlap in the pattern of innervation of the calyx by both MBONa1 and MBONa2 (Fig. 3Bi-iii), as well as in the output region in the ipsilateral (Fig. 3Ci-iii) and contralateral brain (Fig. 3 Civ, Cv, Cvi).

**FIGURE 3.**
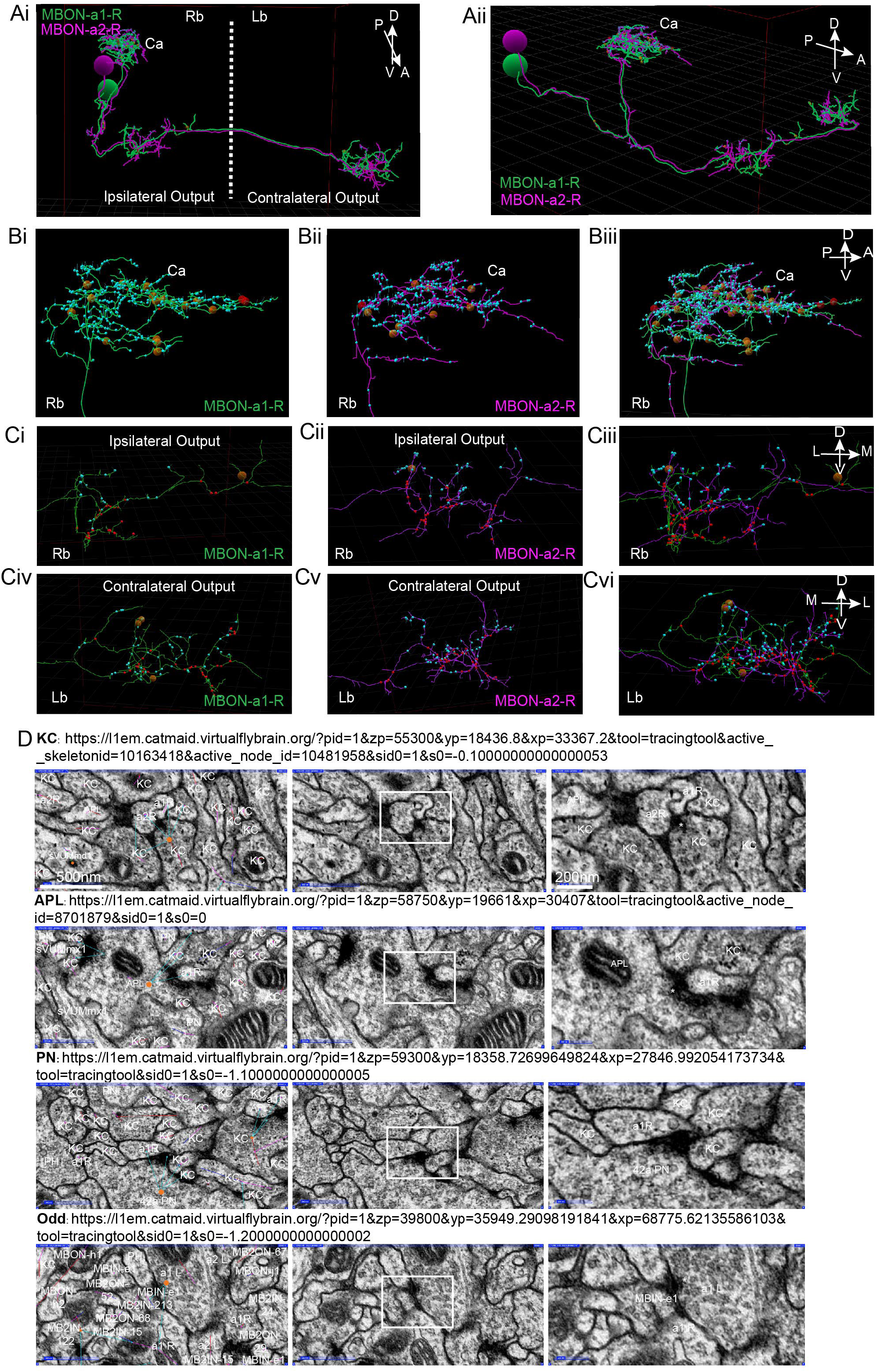
Synapses of MBONa1/a2 visualized by CATMAID in the first instar larva. **(A)** A reconstruction using CATMAID (Eichler et al., 2017) of MBON-a1-R (annotated in CATMAID as MBE7a right) in green and MBON-a2-R (annotated as MBE7b right) in purple, showing their cell bodies (CB), projections in the calyx (Ca), and axonal regions with ipsilateral and contralateral outputs. The axes on the top right corner of the panel indicate the orientation of the neuron, where A is anterior, P is posterior, D is dorsal, V is ventral. **(Ai):** Approximately frontal view of reconstruction. **(Aii):** A more lateral view of the same reconstruction, showing the projections of MBON-a1-R and MBON-a2-R from their posteriorly located cell bodies to the anterior and dorsal parts of the brain. (**B)** A lateral view of a reconstruction of MBON-a1-R **(Bi)** and MBON-a2-R **(Bii)** and their overlap **(Biii)** showing their projections in the right brain calyx. **(C)** Frontal view of the ipsilateral **(Ci-Ciii)** and contralateral **(Civ-Cvi)** outputs around the medial lobes of the MBs of MBON-a1-R **(Ci, Civ)** and MBON-a2-R **(Cii, Cv)** and their overlap **(Ciii, Cvi)**. In all panels, small cyan circles are MBON-a1-R and MBON-a2-R postsynaptic sites, and small red circles are presynaptic sites. The larger brown and red circles are unfinished tracing sites. The axes on the top right corner of panels Biii, Ciii and Cvi apply to each entire row. Images were generated by analysis of neuron tracing using the 3D tool of CATMAID on the publicly available first-instar larval connectome on the Virtual Fly Brain site (https://l1em.catmaid.virtualflybrain.org; Licence CC-BY-SA_4.0). Schneider-Mizell et al., (2016), Saalfeld et al., (2009). **(D)** EM sections of first-instar larva MBONa1/a2 synapses. The top three rows show presynaptic partners of MBON-a1-R (labeled as a1R) in the calyx. Each row shows examples of four common partners: a KC; an APL (labeled as MBE12 on CATMAID); and PN42a. The bottom row of panels shows MBON-a1-L (labeled as MBE7a left on CATMAID), which is presynaptic to contralateral MBON-a1-R in the output regions around the medial lobes. In each row, the left panel shows an EM section with CATMAID annotations; a connector (orange) is placed on the presynaptic neuron and the cyan arrows indicate postsynaptic partners. The middle panel shows the same EM section without the CATMAID annotations. The right panel shows a magnification of the outlined area of each middle panel, showing the characteristic vesicles and T-bar (labeled with an asterisk where present) in presynaptic boutons. The scale bars in the top row apply to all four rows.

Fig. 3D shows representative synapses onto MBONa1/a2 marked in CATMAID: (1) presynaptic terminals of KCs and APL in the calyx, with vesicles and T-bars synapsing on MBONa1/a2, (Fig. 3D top 2 rows); (2) a presynaptic terminal of PN 42a containing vesicles, in the calyx on MBON-a1-R (Fig. 3D, 2nd row from bottom); (3) a presynaptic terminal of MBONa1-L onto MBONa1-R in the output region, suggesting the presence of synapses between MBONa1/a2 there (Fig. 3D, bottom row). In the calyx only a few synapses were from MBONa1/a2 onto KCs, PN13a, and sVUM1s, and no synapses were presynaptic to APL.

### KCs provide the main input to MBONa1/a2 in the calyx

MBONa1/a2 processes in the calyx contain over 305 postsynaptic sites for MBONa1-R and 315 for MBONa2-R, a total of 620 postsynaptic sites in the right calyx. MBONa1/a2 in the left calyx have similar numbers to these. Only 6 presynaptic sites were counted in MBONa1/a2 in the right calyx and none in the left calyx; for MBONa1-R, one is presynaptic to sVUM1 neurons OANa1 and OANa2, a second one to 2 KCs, a third one to a KC and OANa1, and the fourth one to PN13a and a KC. MBON-a2R is presynaptic to 2 KCs, and presynaptic in a divergent synapse to an unidentified neuron “place holder neuron” and 2 KCs. MBONa1/a2 had no presynapses onto APL. Therefore, MBONa1/a2 are mainly postsynaptic in the calyx (Supplementary Table 1).

KCs provide the main input into MBONa1/a2 in the calyx, with 314 presynaptic sites on MBONa1-R out of 349, about 90% of its total input in the calyx. Taking into account that there are 110 KCs on the left brain and 113 on the right in the first instar connectome (Eichler et al. 2017), it is possible that nearly all KCs have presynaptic contacts with MBONa1-R (Fig. 4). The remaining presynaptic inputs onto MBONa1-R are from sVUM1 neurons with 17 inputs (4.9%), APL with 14 inputs (4.0%), and only 4 inputs (1.1%) were from PNs. The proportions of presynaptic partner inputs onto each of the other MBONa1/a2 were similar to those on MBON-a1-R (Fig. 4). There was no preference for synapses with particular types of KC; KC39, KC54, and KC77 have three to four dendrites that are spread out, while dendrites of KC34, KC25 and KC49 KC22 were localized to a subregion of the calyx for the right brain. Some KCs provide input to both MBONa1/a2 neurons in the right brain.

**FIGURE 4.**
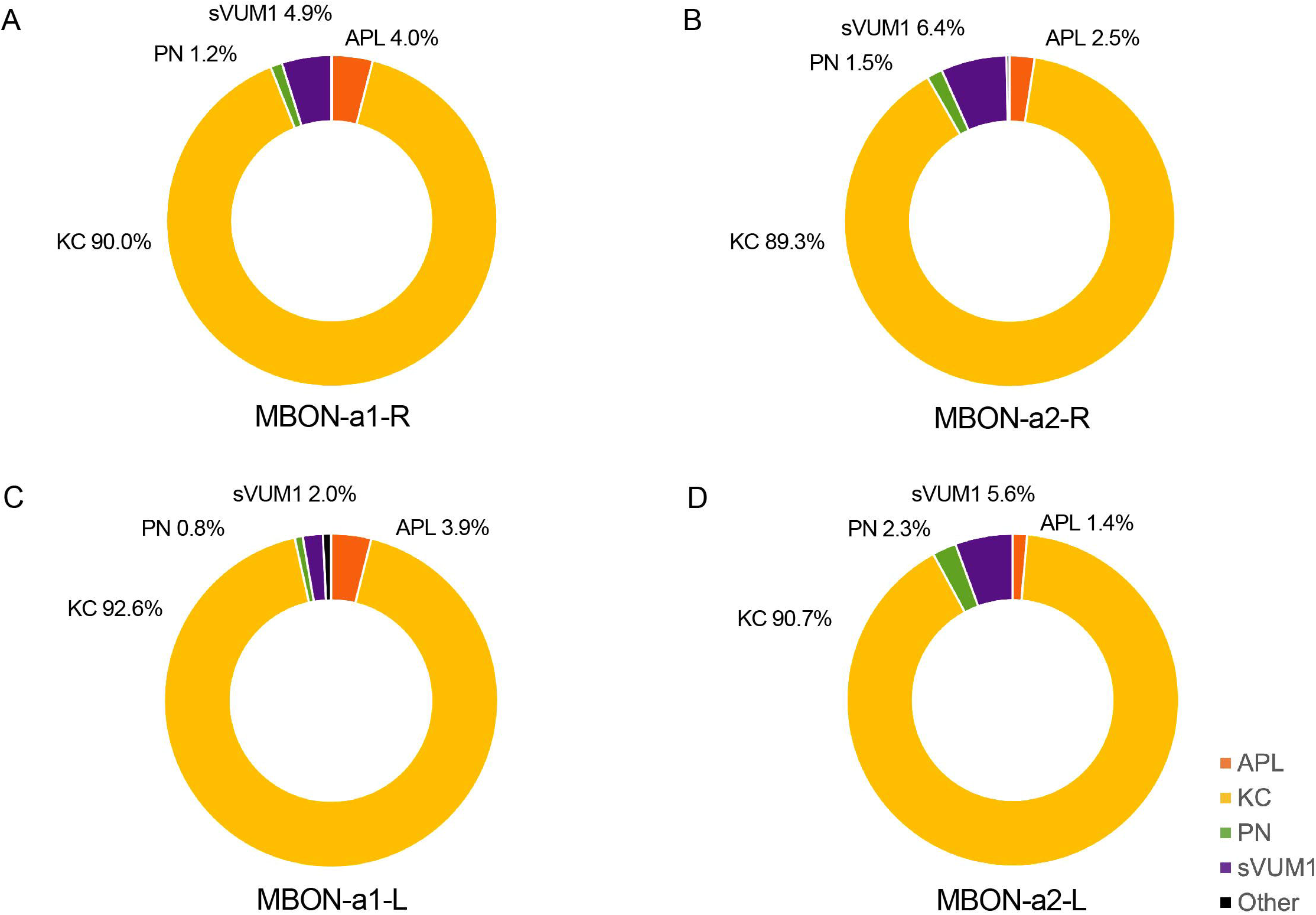
The types of neurons presynaptic to MBONa1/a2 in the calyx. This figure shows the proportion of KCs, APL, PN, sVUM1 neurons and other neurons that provide input to MBONa1/a2 in the calyx. These were calculated from CATMAID data by dividing the number of presynaptic inputs to MBONa1/a2 from each neuron type by the total number of annotated inputs of each MBONa1/a2 neuron in the calyx (See Materials and Methods). APLs are shown in orange, KCs in yellow, PNs in green, sVUM1s in purple and other types of neurons in black. Individual panels show the types of calyx presynaptic partners of (**A**) MBON-a1-R, (**B**) MBON-a2-R, (**C)** MBON-a1-L and (**D**) MBON-a2-L.

### PN, sVUM1s, APL, inputs to MBONa1/a2 in the calyx

The second main inputs to MBONa1/a2 are from the two sVUM1 neurons innervating the calyx (OANa1/a2), with 5% of total inputs onto MBON-a1-R. APL inputs are 4% of the total inputs to MBONa1-R, which is significant considering that there is only a single larval APL in each side of the brain. APL in the right brain has 14 and 8 presynaptic sites on MBONa1-R and MBONa2-R, respectively, while APL in the left brain has 10 and 6 presynaptic sites and MBONa1-L and MBONa2-L, respectively.

The number of inputs onto MBONa1/a2 neurons from PNs was only around 1% for MBON-a1-R, 4 synapses out of 349. These numbers are surprisingly low, considering that there are at least 21 PN inputs in the calyx, and raises the question of whether these connections are functionally significant. Also arguing against significant PN-MBONa1/a2 connectivity, the EM sections of PN-MBONa1/a2 synapses did not meet most of the synapse criteria, e.g. a PN42a synapse showed no T-bar, postsynaptic density or synaptic cleft (Fig. 3D, second row from bottom), supporting the doubts on the roles of PN-MBONa1/a2 contacts in the calyx.

### Mix of pre and post synapses in the Output of MBONa1/a2

On the other hand, the output region of the MBONa1/a2around the MB medial lobe contained a mix of postsynaptic and presynaptic sites, segregated in axonal branches that were compartmentalized into either mainly presynaptic or postsynaptic sites (Fig. 3 Ci to Cvi). The number of presynaptic sites for MBON-a1-R in the right brain was 35, and 33 for MBON-a2-R, a total of 68; the equivalent numbers of postsynaptic sites were 20 for MBONa1-R and 36 for MBONa2-R, a total of 56. Similar numbers were counted in the contralateral left brain (Supplementary Table 2), where MBON-a1 and MBON-a2 had similar numbers of synapses (Supplementary Fig. 2). Therefore, the output regions of MBON-a1 and MBON-a2 contain comparable numbers of presynaptic and postsynaptic regions, in the ipsilateral and contralateral output regions. Interestingly, MBONa1/a2 target many MBONs and MB inputs in this area (Supplementary Fig. 3).

Some reciprocal synapses between MBONa1 and MBONa2 were also found in the output region. As shown in Fig. 3D, bottom panel, the synapse from MBON-a1-L to MBON-a1-R shows a visible T-bar and vesicles in MBON-a1-L. All four MBON-a1/-a2 neurons have synaptic connections among each other, numbering between 1 and 8, and no differences were detected between ipsilateral and contralateral partners (Supplementary Table 3). There were no annotated synapses between MBONa1/MBONa2 neurons in the calyx. Only 11 presynaptic sites onto KCs were annotated in the lobes, indicating that MBONa1/a2 do not provide significant output to KCs around the medial lobes.

### Odor concentration dependence of MBONa1/a2 neuron responses

Despite the lack of strong GRASP or anatomical evidence for PN-MBONa1/a2 contacts, Odd neurons have been implicated in concentration-dependent behavioral responses to odors (Slater et al., 2015). We therefore analyzed the concentration-dependence of MBONa1/a2 responses to odors, by expressing the genetically encoded calcium indicator *jRCaMP1b* (Dana et al., 2016) in MBONa1/a2 using the split-GAL4 line *64FO7-p65.AD* (X); *R57C10-GAL4DBD (III)* (Fig. 1), and recorded calcium responses to different concentrations of ethylacetate (EA). For comparison, we also assayed odor responses using *jRCaMP1b* expressed in KCs, known to have relatively concentration-invariant responses (Stopfer at al., 2003).

Odor-evoked activity was observed throughout MBONa1/a2, including in the calyx and output regions around the ML. In the calyx, activity was predominantly localized to regions around glomeruli (Fig. 5A, left panel); the latter interpretation is consistent with the interglomerular localization of MBONa1/a2 processes (Fig. 1B). Some activity was also observed in the non-glomerular calyx core, and the dendritic shaft that leaves the calyx (Fig. 5A, arrow). The odor-evoked response pattern of MBONa1/a2 in the calyx is similar to the activity of KCs in response to EA, where a few glomeruli are activated by EA (Fig. 5A, right panel), as expected from our previous observation that odors are represented in a spatially localized pattern in the calyx (Masuda-Nakagawa et al., 2009). Since *MB247-GAL4* is a strong driver that labels most KCs, of which there are about 2100 at larval wandering stage (Technau and Heisenberg, 1982), and KCs show a dense pattern of innervation of the calyx in glomerular and non-glomerular regions, a strong baseline fluorescence was observed.

**FIGURE 5.**
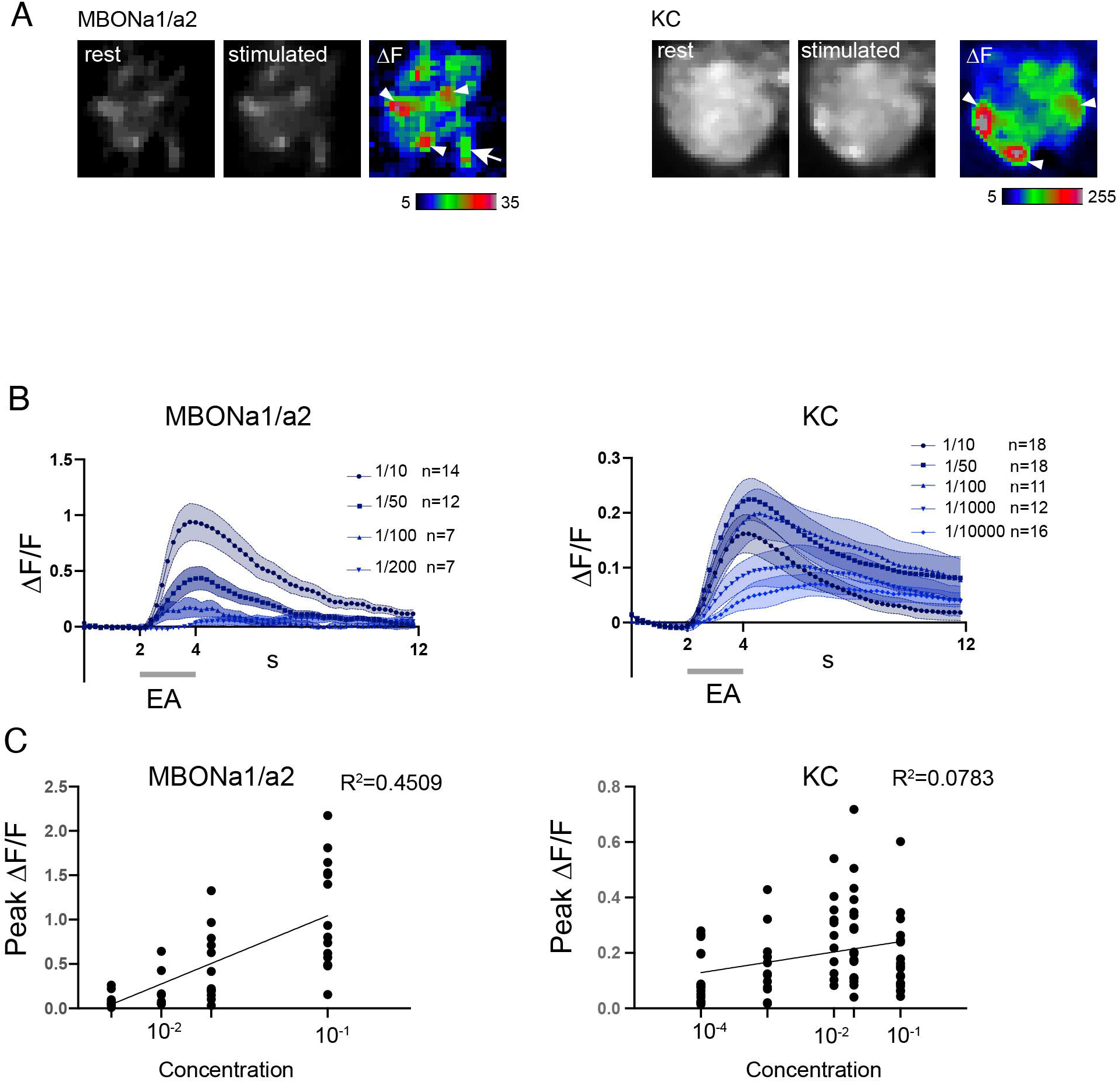
MBONa1/a2 responses to changes in concentration of ethyl acetate are more graded than KC responses. **(A)** Odor-evoked activity in MBONa1/a2 expressing *UAS-jRCamP1b* using *64FO7-p65.AD (X); R57C10-GAL4DBD (III)* (left panels) and in KCs expressing *MB247-GAL4* (right panels) in response to a 2-second pulse of ethylacetate (EA) diluted 10-fold in mineral oil. Resting and peak fluorescence are shown in greyscale, and ΔF in pseudocolor. White arrowheads show areas of elevated response levels corresponding to glomeruli in the KC panels, and which may represent areas adjacent to specific glomeruli in the MBONa1/a2 panels. **(B)** Time course of odor-evoked responses to different concentrations of ethylacetate (EA) in MBONa1/a2 or KCs. A 2-second odor pulse is indicated by a grey bar. Graphs show mean ± SEM. A three-point moving average was used to smoothen responses. **C**. Graphs of peak ΔF/F against log (concentration of ethyl acetate), with each individual data point shown. Each line represents the calculated linear regression for MBONa1/a2: Y= 0.7696X + 1.814, R^2^ = 0.4509, slope significantly different from 0 (two-tailed P<0.0001****) with the Pearson test. Linear regression for KCs: Y= 0.03727X + 0.2779, R^2^ = 0.07827, slope significantly different from 0 (two-tailed P=0.0151*) with the Pearson test.

The MBONa1/a2 odor response began at the onset of the odor pulse, except for the lowest dilution at 1/200 where the small increase started only at the end of the odor pulse (Fig. 5B). The time course of KC responses showed a similar temporal pattern at the onset of the odor pulse, but the ΔF/F values were lower (Fig. 5C). This could potentially be an effect of the high baseline fluorescence observed in the calyx using the *MB247-GAL4* driver.

MBONa1/a2 responded to high concentrations of odors, e.g 1/10 dilution of EA, but not to 1/200 and very weakly to 1/100 dilutions of EA (Fig. 5B). The peak ΔF/F gradually decreased with increasing odor dilution, with a 1/100 dilution of EA giving a mean response around 21% of that of a 1/10 dilution, and a 1/200 dilution only 10%, and not statistically different from zero (P=0.194, two-tailed t-test). The number of preps without an odor response increased from 0% at 1/10 dilution, to 1-2 at 1/50 and 1/100 dilutions, and at 1/200 dilution only 2 preps out of 7 showed a detectable odor response. These findings suggest that MBONa1/a2 responses have a narrow dynamic range, but are strongly concentration-dependent (Fig. 5C), with a slope that is significantly different to zero in a linear regression analysis.

In contrast, the odor-evoked responses of KCs were less concentration-dependent. For all odor dilutions, KC responses closely followed odor onset. At all EA dilutions except for 1/10000, a clear odor-evoked response was observed, and 25% of preparations were non-responsive at 1/10000. KC responses were less concentration-dependent than those of MBONa1/a2, showing similar peak ΔF/F at EA dilutions of 1/10, 1/50, 1/100, 1/1000, and only a shallow slope in the regression analysis (Fig. 5C).

### Optogenetic stimulation of OA neurons does not have a detectable effect on odor-evoked responses in MBONa1/a2

The calyx of the larval MBs is densely innervated by the terminals of octopaminergic neurons. We have previously shown that MBONa1/a2 labeled by *OK263-GAL4* have potential synaptic contacts with *Tdc2*-expressing octopaminergic neurons innervating the calyx (Wong, Wan et al., 2021). Since octopamine is structurally and functionally similar to noradrenaline, and is a positive modulator of circuit activity (Roeder, 2005, Strother et al., 2018), we tested whether activity of *Tdc2*-OA neurons could potentially enhance odor responses in MBONa1/a2. We measured MBONa1/a2 odor-evoked responses using *52E12-GAL4* and *UAS-JRCaMP1b*, in the presence of a *LexAop-ChR2-XXL* construct and the OA driver line *Tdc2-LexA*, and in control larvae lacking either *LexAop-ChR2-XXL* or *Tdc2-LexA*. We measured responses in individual calyces, and the output region of MBONa1/a2 to (sequentially) (i) an odor pulse (“odor-only”); (ii) an odor pulse delivered immediately after the end of a blue light pulse (“light+odor”). In one set of experiments, procedures (i) and (ii) were followed by (iii) a pulse of blue light to activate ChR2-XXL if expressed (“light-only”).

In larvae of the experimental genotype, odor-evoked responses in MBONa1/a2 in the calyx, were localized to a few high-activity regions that resemble interglomerular spaces or regions bordering glomeruli, consistent with the dense innervation of MBONa1/a2 ramifying throughout the non-glomerular regions of the calyx (Fig. 6A). In the output region in the vicinity of the ML (Fig. 1C), an odor-evoked response was observed all along the axonal processes of MBONa1/a2 (Fig. 6A, right panels). The time course of the odor-evoked response (ΔF/F) showed an increase in response after the onset of the odor pulse for “Odor only”, whereas Light+Odor showed an increase starting with the light pulse, suggesting that there is an odor-independent effect of light activation of ChR2-XXL on the activity of MBONa1/a2 (Fig. 6B). Comparisons between “Odor only” odor-evoked peak ΔF/F with and without prior light pulses showed that the difference is significant (Fig. 6B, bottom panel).

**FIGURE 6.**
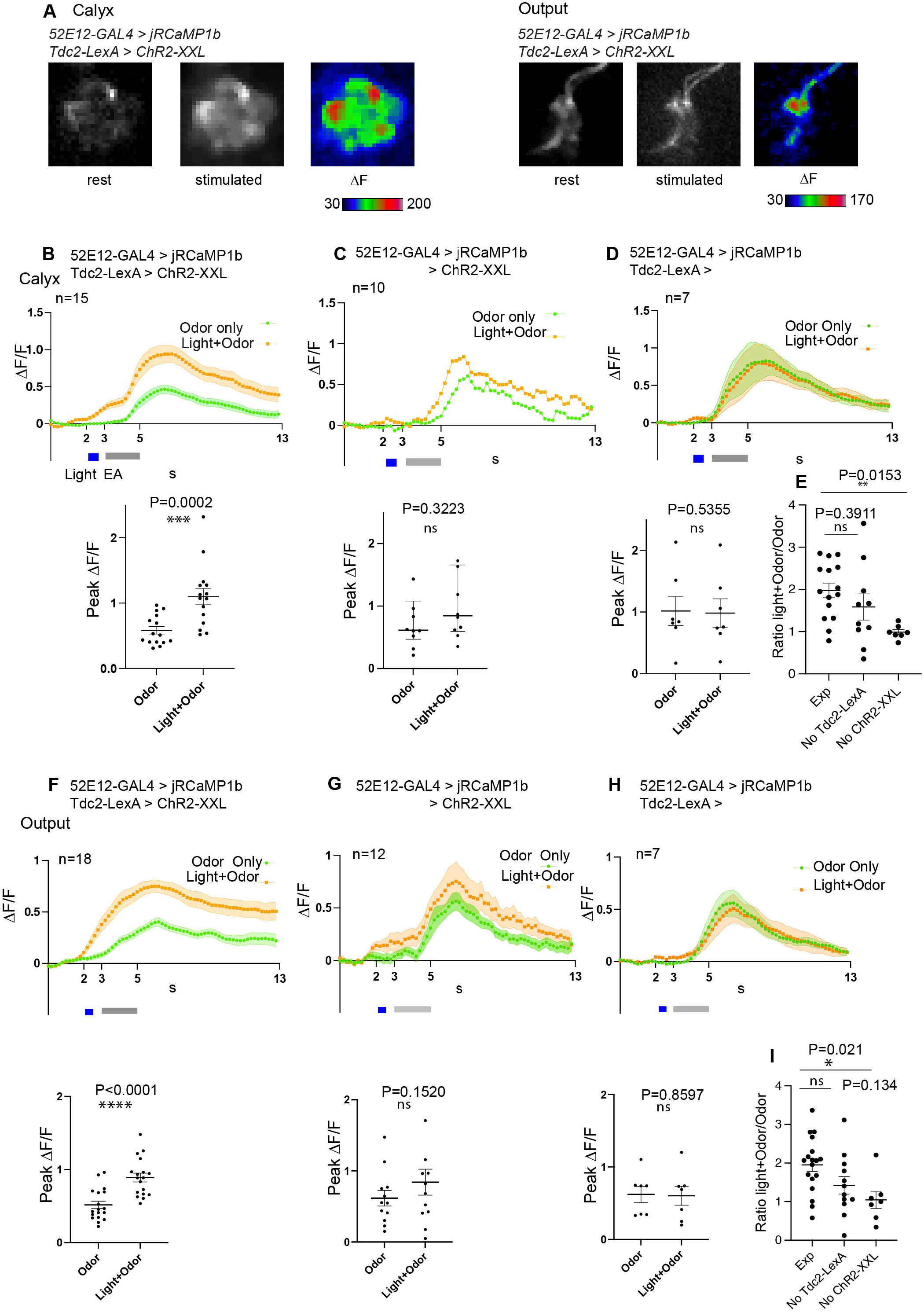
Activation of Tdc2-expressing neurons has only an additive effect on odor-evoked activity of MBONa1/a2 in the calyx and axonal region. **(A)** Odor-evoked activity in MBONa1/a2 expressing *UAS-jRCamP1b* under the control of *52E12-GAL4*, and carrying *Tdc2-LexA* driving *LeAop-ChR2-XXL*, to a 2s pulse of ethylacetate (EA) diluted 10-fold in mineral oil, after previous 561-nm light exposure to activate ChR2-XXL. Responses in the calyx (left panels) or the axonal output area L2, area around ML as shown in Fig 1C (right panels) are shown. Projections of confocal images in grayscale show fluorescence at rest before odor stimulation, and after stimulation, and ΔF shown in color. **(B)** Time course of odor responses of MBONa1/a2 to a 2s pulse of EA in the same brains either before or after light stimulation to activate ChR2-XXL in *Tdc2-LexA*-expressing neurons, in the calyx. Genotype is as in A. The upper graph shows a time course with mean ± SEM of ΔF/F, and a three-point moving average was used to smoothen responses. The grey bar shows the EA pulse, and the light pulse (for Light+Odor) is shown in blue. The scatter graph shows peak odor-evoked responses, without light stimulation (Odor) or following optogenetic stimulation (Light + Odor). Data from two sets of crosses were merged; a t-test showed no significant difference between the peak ΔF/F values from the 2 crosses. **(C)** Time courses of MBONa1/a2 odor responses in larvae of the same genotype as A, but lacking *Tdc2-LexA*, stimulated by odor only or odor-evoked response after previous light pulse as in B. Normality for the peak ΔF/F responses could not be proven, therefore the time course shows median values, and the scatter graph shows peak ΔF/F responses with median and interquartile range. Data from two sets of crosses were merged; a Mann-Whitney test showed no significant difference between the peak ΔF/F values from the 2 crosses. **(D)** Time courses of MBONa1/a2 responses, and peak ΔF/F responses, from larvae of the same genotype as A but lacking the *LexAop-ChR2-XXL* reporter, stimulated by odor only or odor-evoked response after previous light pulses as in B. **(E)** Comparisons of the ratio of peak ΔF/F in Light + Odor conditions to that in Odor conditions in the calyx, between larvae of the same genotype as A and B, and controls lacking either *Tdc2-LexA* (as in C) or *LexAop-Chr2-XXL* (as in D). Comparisons by an ordinary one-way ANOVA test followed by Tukey multiple comparison tests are shown. **(F-I)** Responses and comparisons as in panels B-E respectively, but measured in the MBONa1/a2 axonal output region around the ML. Peak ΔF/F values in G met a normality test and therefore data are plotted as mean ± SEM, and compared using a t-test. In all quantitation panels, data were tested for normality using a Kolmogorov-Smirnov test. In Panels B-D and F-H, time courses of normally distributed data are shown as mean ± SEM. For non-normally distributed data (C), only median is shown. Normally distributed peak ΔF/F responses are presented in scatter graphs as larval datapoints, mean ± SEM, and compared using a two-tailed paired t-test. Non-normally distributed peak ΔF/F responses are presented in scatter plots as larval datapoints, median ± interquartile range, and compared using a two-tailed Wilcoxon test. Occasional outlier datapoints are omitted from the scatter graphs, but included in all statistical analyses. For E and I, normality within each data set was confirmed using a Kolmogorov-Smirnov test. The graph shows individual larval datapoints, mean ± SEM.

To directly compare the effect of specific activation of ChR2-XXL in Tdc2-neurons with responses in controls, we compared the ratio of “Light+Odor” to “Odor only” responses, between larvae of the experimental genotype, and larvae lacking the expression of either *Tdc2-LexA* (hence potentially expressing *ChR2-XXL* non-specifically), or larvae lacking the *ChR2-XXL* (Fig. 6E). In agreement with the measurements in Fig. 6B-D, this ratio was higher in the experimental genotype compared to both controls, but only the effect of lacking ChR2-XXL was significant; the effect of lacking *Tdc2-LexA* was not significant (Fig. 6E). The results suggests that there is an effect of light on MBONa1/a2 responses that may be caused by non-specific expression of ChR2-XXL. In the output region, where we are measuring axonal or presynaptic responses (Fig. 1D, Fig. 6A, right panels), our findings (Fig. 6F-I) were consistent with the those in the calyx, again suggesting an effect of light on ChR2-XXL activation that is non-specific to *Tdc2* neuron activity.

The theoretical sum of Odor-only response and the Light-only response closely matched that of the odor-evoked response with prior light pulses, and showed no significant difference in their peak (ΔF/F) responses, suggesting that the effect of a prior light pulse on the odor response of the experimental genotype can be accounted for by an additive effect of light and odor (Supplementary Fig. 4, Ai-Aiii), and is not due to regulation of MBONa1/a2 responses by *Tdc2-LexA* neurons. We reached a similar conclusion for MBONa1/a2 responses in the output region, albeit with a lower response to light-only stimulation (Supplementary Fig. 4, Bi-iii). Taken together, we therefore find no evidence for an effect of sVUM1 neuron activity on odor-evoked MBONa1/a2 activity under the conditions tested here.

### Optogenetic activation of MBONa1/a2 neurons impairs behavioral odor discrimination learning

MBONa1/a2 responds only to higher concentrations of odors (Fig. 5). We therefore tested the hypothesis that MBONa1/a2 affect discrimination learning in an odor concentration-dependent manner, by optogenetic activation of MBONa1/a2 using the long-wavelength-absorbing channelrhodopsin, CsChrimson (Klapoetke et al., 2014) driven by the split line *64F07-p65.AD* (X); *R57C10-GAL4DBD (III)*. As controls we used the progeny of the *Canton S* line crossed to the reporter CsChrimson. We measured learning scores after conditioning with amber light (to activate CsChrimson if expressed) or blue light as control, at two different concentrations of EA and pentyl acetate (PA): a higher concentration with dilutions of EA2000:PA500 and a lower concentration with dilutions of EA4000:PA1000.

Activation of MBONA1/a2 neurons in amber light during conditioning led to a lowered learning score compared to blue light at both odorant concentrations tested (Figure 7). A 2-way ANOVA test on this genotype showed a significant effect of amber light (P = 0.0193), and no significant effect of concentration (P > 0.4). While conditioning under amber light appeared to have a larger effect at the higher odorant concentration than at the lower one, ANOVA showed no significant interaction between light and concentration (P > 0.6), suggesting that the effect of light (and CsChrimson activation in MBONa1/a2) was statistically indistinguishable across both concentrations. A similar analysis of the control cross showed similar learning scores to the experimental cross under blue light, but in contrast to the experimental cross, there was no effect of amber light (P > 0.6). As in the experimental cross, there was no effect of concentration in the control cross (P > 0.05). We therefore conclude that activation of MBONa1/a2 impairs olfactory discrimination reward learning in larvae, but that there is not a statistically detectable dependence of this effect on odor concentration.

**FIGURE 7.**
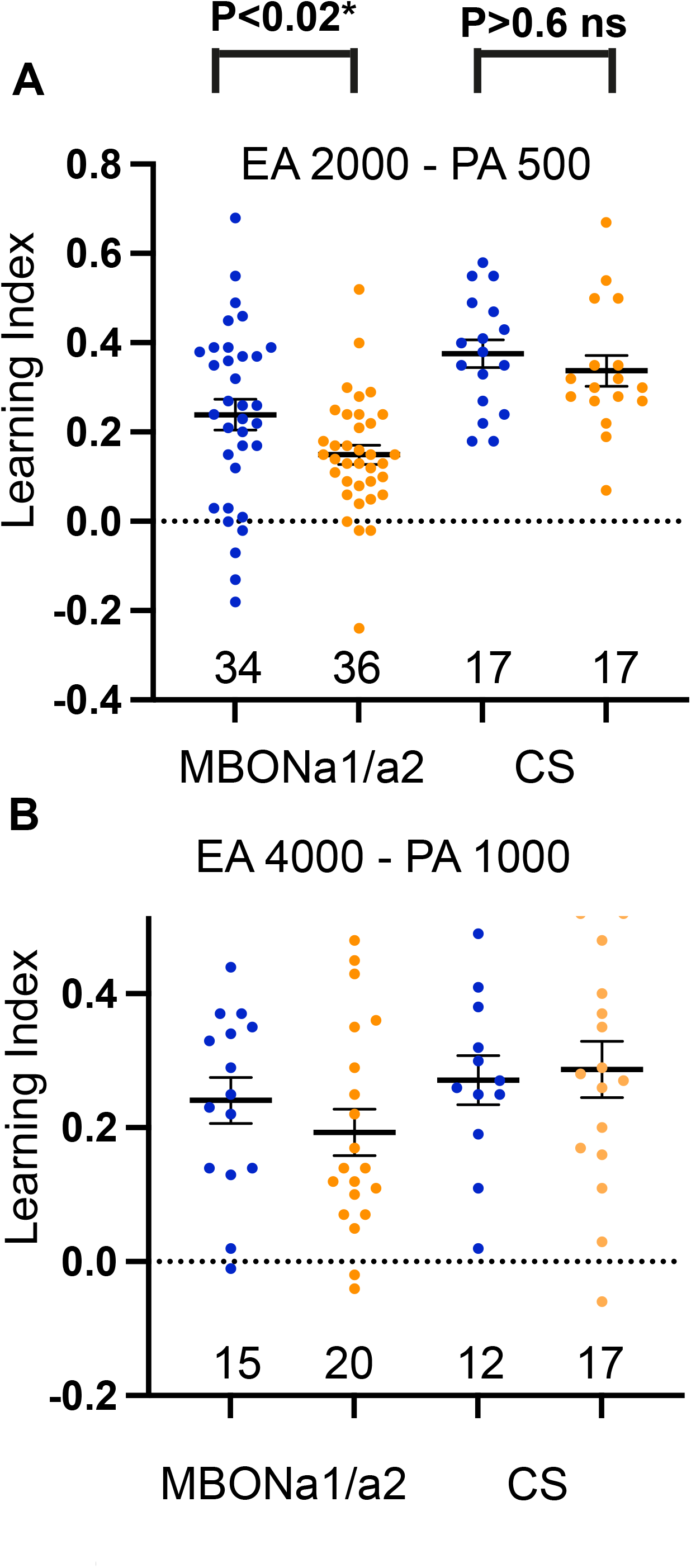
Activation of MBONa1/a2 impairs odor discrimination learning. **(A)** Odor discrimination learning using EA at 1/2000 and PA at 1/500 dilution. Blue dots and amber dots indicate conditioning under blue light, or under amber light, which activates CsChrimson. Genotypes were the progeny of UAS-CsChrimson crossed to either the split-GAL4 line *64FO7-p65.AD* (X); *R57C10-GAL4DBD* (III) (“MBONa1/a2”) or to CantonS controls (“CS”). (**B)** Same as A, using EA at 1/4000, and PA at 1/1000 dilution. Two-way ANOVA (light and concentration) shows a significant effect of amber light on learning across concentrations for MBONa1/a2 activation (P=0.0193), but no significant effect on learning scores in CS controls lacking the split driver (P=0.69). Interaction between light and concentration was not significant in ANOVA (P = 0.64), showing that the effect of MBONa1/a2 activation on learning was not concentration-dependent within the range of concentrations tested.

## Discussion

### MBONa1/a2 show major contacts with KCs, APL, and sVUM1s but not PNs

The larval calyx is organized in about 34 glomeruli, sites of PN cholinergic terminal boutons synapsing with KC dendrites that are visualized by anti-DLG as the shape of a bouquet of round structures around a central core (Masuda-Nakagawa et al., 2005). MBONa1/a2 arborize in the core of the calyx and interglomerular space, with processes abutting glomeruli but clearly avoiding their interior, suggesting that synapses onto MBONa1/a2 occur at non-glomerular space, suggesting that they might not receive direct PN input.

Contrary to our expectations based on Slater et al., (2015), who showed GRASP signals between PNs and Odd neurons in the calyx, GRASP signals were rare or atypical between PNs and the MBONa1 or MBONa2 labeled *by 68B12-GAL4*. Long axonal tracts bordering the inner surface of a few glomeruli were observed labeled with GFP. There was no co-localization of GRASP with OA, suggesting that these tracts were not likely to be non-glomerular synaptic sites of PNs onto sVUMs and MBONa1/a2. The possibility exists that MBONa1/a2 would make synapses with PNs in the core of the calyx, where PN axons are also traversing, however, GRASP signal between MBONa1/a2 and PNs were not observed in the core of the calyx. Our evidence favors the view that PNs do not provide direct input onto MBONa1/a2, suggesting that olfactory input is not delivered directly onto MBONa1/a2 by the olfactory PNs.

We have shown previously that MBONa1/a2 labeled by *OK263-GAL4* have many contacts with OA neurons at interglomerular sites (Wong, Wan et al., 2021, Fig. 3D). Here we have shown that KCs also have contact sites with the MBONa1 or a2 labeled by *68B12-GAL4* in regions neighboring glomeruli. GRASP signals between KCs and MBONa1/a2 resemble a mesh rather than the typical dot-like signals. This might be caused by the density of KC processes in the core of the calyx and limited space between glomeruli making it easier for MBONa1/a2 processes to become in close proximity to KC processes. Nevertheless, typical GRASP signals with high GFP intensity were seen near OA labeling, suggesting proximity to synapses between KCs and sVUM1 neurons (Fig. 2B), a feature observed in previous EM analysis, showing divergent synapses of sVUM1 neurons onto MBONa1/a2, APL and KCs (Wong, Wan et al., 2020, Fig. 5E bottom three panels). KC dendrites are known to have presynaptic sites at a distance from the dendritic region containing the synapses between PNs and KCs, (Christiansen et al., 2011), therefore KC contacts in the calyx core or interglomerular space could be KC presynaptic sites onto MBONa1/a2.

Contacts between the larval APL and the single MBONa1/a2 neuron labeled by *68B12-GAL4* formed characteristic dot-like GRASP signals, that overlapped with GABA often at one end of a large GABAergic APL bouton (Fig. 2Biii). Not all GABA termini showed GRASP signals, and GRASP signals were in the calyx core and interglomerular space. Since APL is presynaptic in the calyx (Wong, Wan et al. 2020), this suggests that APL synapses onto MBONa1/a2.

We found that MBONa1/a2 processes have a clear polarity (Fig. 1D): in the calyx processes are mainly postsynaptic, labeled by the postsynaptic marker DenMark::mCherry, and only a few puncta labeled by Syt::GFP or n-Syb::GFP (Supplementary Fig. 1D), whereas the output region surrounding the medial lobe is presynaptic and labeled by Syt::GFP and nSyb::GFP, suggesting that MBONa1/a2 receive predominantly inputs at the calyx. This suggests that APL, sVUM1s, and even KCs are likely presynaptic to MBONa1/a2 in the calyx, suggesting a role for MBONa1/a2 as integrators of multiple inputs, rather that relay neurons of olfactory input.

### Comparison with the first instar connectome

The first instar (6-hr) larva connectome provides a comprehensive map of synaptic connections (Eichler et al., 2017). However, neurons can be immature at this stage, and therefore we compared the annotated connections with those that we found in the more mature 3rd instar larvae. MBONa1/a2, known also as MBE7a/b in the connectome, shows a similar morphology to MBONa1/a2 in the 3rd instar; 2 neurons have been annotated with similar and spatially overlapping projections to the calyx, ipsilateral medial lobe and contralateral medial lobe. MBONa1/a2 processes in the first-instar calyx were postsynaptic: in the right brain, only 6 synapses were presynaptic compared to more than 600 postsynaptic sites, in agreement with our observations in 3rd instar larvae.

We have previously shown that at the EM level, sVUM1 neurons have divergent synapses onto MBONa1/a2, KCs and APL; and that these synapses are surrounded by many KC processes (Wong, Wan et al. 2021, Fig 5E bottom three panels). Here we found that synapses of KCs on MBONa1/a2 are also divergent, synapsing onto MBONa1, MBON a2, and KCs (Fig. 3D, top panel); APL also shows divergent synapses onto MBONa1 and a KC. Therefore, in the calyx synapses between the different types of calyx neurons, KCs, APL, sVUM1s, and MBONa1/a2 appear to be co-localized. This is consistent with our GRASP analysis between KCs and MBONa1/a2, where GRASP signal co-localizes with OA labeling in the calyx, suggesting the presence of synaptically rich regions localized to regions that neighbor glomeruli or in the core of the calyx.

Our CATMAID analysis shows that only a few PNs synapse onto MBONa1/a2. Since there are at least 21 olfactory PNs, it is unlikely that PNs are a major input into MBONa1/a2. This is consistent with our GRASP results in which GRASP signals between PNs and MBONa1/a2 labeled tracts with no distinct punctate localization, suggesting that the signal on tracts may not be synaptic contacts as interpreted by Slater et al., 2015 (Fig 3A).

We found that KCs are the major input to MBONa1/a2 in the calyx. KCs form more than 600 synapses onto MBONa1/a2 in the right brain calyx, which are 90% of total MBONa1/a2 calyx inputs, and considering that there are 223 KCs at this early stage in the brain, summing up left and right brain KC numbers, potentially all KCs could input into MBONa1/a2. sVUM1 and APL also have inputs into MBONa1/a2, suggesting that MBONa1/a2 can be activated by KCs, but also integrate calyx inhibitory and modulatory input.

A difference from the 3rd instar larvae was found in the output region of first-instar MBONa1/a2, where synapses were a mix of pre- and post -synaptic sites without a defined polarity: 68 presynaptic sites and 56 postsynaptic sites in the right brain. However, there was a regional segregation of these synapses, with branches that were exclusively presynaptic or postsynaptic. This is in contrast to the 3rd instar output region which appears exclusively presynaptic using presynaptic and dendritic markers (Fig. 1D). This might be due to a developmental issue, if the postsynaptic projections in the medial lobe contain immature synapses that are subsequently removed or pruned. Immature synapses are common at this stage; for example for KCs, nearly 65% are immature, defined by lack of dendrites, or tiny dendrites and processes with endings typical of growth cones (Eichler et al., 2017). The output region had synapses between MBONa1/a2 in CATMAID; however, presynaptic vesicles were ambiguous and there were only a small number of synapses (Supplementary Table 3), questioning whether these were true synapses. However, this observation raises the possibility that MBONa1/a2 may provide bidirectional input to each other.

### Concentration dependence of MBONa1/a2

Our anatomical studies show that MBONa1/a2 neurons ramify extensively throughout the calyx core and interglomerular space and receive few direct PN inputs, casting doubt on whether they might be activated by odor input. However, specific expression of jRCAMP1b in MBONa1/a2 using the split line *64FO7-p65.AD (X); R57C10-GAL4DBD (III)* showed odor-evoked response in the postsynaptic dendrites in the calyx at localized regions (Fig. 5, 6). Moreover, the response was concentration-dependent, showing graded responses to increasing concentrations of odor. The dynamic range was narrow, lying between a 10-fold and 200-fold dilutions, and the latter concentration gave only a negligible response. In sharp contrast, KCs responded to odors at a dilution as low as 10^−4^. Also in adult flies, KCs respond to odorant dilutions of 10^−5^ to 10^−2^ (Wang et al., 2004). Therefore, MBONa1/a2 have a high threshold of activation. Their extensive ramification in the calyx predicts a high capacitance that could rise the threshold of firing, compared to KCs that innervate only a few to 5 glomeruli in the larva (Masuda-Nakagawa et al., 2005). This is the first time that the specific concentration-dependence of MBONa1/a2 is reported; Slater et al., (2015) analyzed concentration-dependence for the apple cider vinegar, ACV, in *Odd-GAL4*-expressing cell bodies, that they reported to label 8 neurons including 3 innervating the calyx, only within a relatively high concentration range of 10-fold to 20-fold dilution. The concentration dependence of MBONa1/a2 reported here, in light of their anatomical organization that shows no regional arborization of dendrites in the calyx, suggests that they have a role in odor intensity coding, as opposed to KCs that work in a combinatorial mechanism that allows odor discrimination and encoding of many odors.

CATMAID and GRASP analysis in 3rd instar larvae are consistent with MBONa1/a2 receiving their major input from KCs, sVUM1s, and APL neurons innervating the calyx. This implies that the source of olfactory input onto MBONa1/a2 might be an indirect activation of MBONa1/a2 mediated by other calyx neurons; for example KC dendrites, and APL terminals in the calyx (Masuda-Nakagawa et al., 2014) are activated by odors. KCs are known to possess presynapses at the dendrites in the calyx (Christiansen et al., 2011) and would be the most obvious candidate to activate MBONa1/a2 dendrites. The odor-evoked response by KCs in the calyx, showed activation of a few glomeruli, consistent with one odor activating a few olfactory receptors that act in a combinatorial manner (Hallem et al., 2004). In flies it has also been shown that the EA-evoked odor response in KCs localizes to particular calycal areas (Wan et al., 2004). The odor-evoked response of MBONa1/a2 in the calyx also showed preferential localized activation, as opposed to whole activation of the dendritic processes in the calyx, similar to the pattern of localized glomerular activation observed in the calyx by the odor-evoked activation of KCs, suggesting that MBONa1/a2 dendritic processes in the vicinity of KC processes activated in glomeruli by PN input, might receive input from KCs. The picture that emerges is that MBONa1/a2 are postsynaptic to KCs, generating an output channel that encodes the intensity of olfactory input indirectly.

### State dependent signals and MBONa1/a2

Octopaminergic neurons are positive neuromodulators of circuit function, for example in the motion vision system in the fly, they provide input to motion detection neurons and are involved in their temporal tuning, by increasing the excitability of medulla input neurons (Strother et al., 2017). We have previously shown that activation of sVUM1s OA neurons innervating the calyx, impair the behavioral discrimination of sensory stimuli (Wong, Wan et al., 2021), suggesting a role of calyx-innervating OA neurons in modulating calyx circuit activity. To test whether sVUM1 neurons in the calyx would modulate the excitability of MBONa1/a2, we activated Tdc2 neurons, by optogenetic stimulation of ChR2-XXL. Localized odor-evoked responses were observed in both the calyx and output region of MBONa1/a2 upon prior optogenetic activation of sVUM1 neurons, similar to an odor response in the specific split line (Fig. 5A). The time course of odor plus light response in the calyx showed an earlier rise immediately after the light pulse and was also longer lasting than the odor only response which return to baseline after 13 seconds, showing that light alone has an effect on MBONa1/a2 activity. On the other hand in the output region, the odor response did not return to baseline after 13s, and the light plus odor response still remained high after 13 seconds. The difference between the response in the calyx and the output response could be mediated by the larval APL, a feedback neuron, that would shut down the response in the calyx, since MBONa1/a2 receive significant input from APL in the calyx, also shown by GRASP signals between MBONa1/a2 and APL.

When comparing the theoretical sum of the light only response and odor only response to the experimental odor-evoked response with prior light stimulation, there was no enhancement of the odor-evoked response in MBONa1/a2 in the calyx nor the output region, but only an additive effect of light stimulation. It is interesting to note that the light only response of MBONa1/a2 in the calyx shows an increase in response, while the light only response in the output region showed no increase by light alone, suggesting that the effect of light is specific to the activation of ChR2-XXL upstream of the MBONa1/a2 in the calyx. Our controls without driver Tdc-GAL4 showed no statistically significant difference between odor-evoked response with or without prior light stimulation, suggesting that at least some effect could originate in Tdc2 activation. Therefore, activation of sVUM1 neurons did not have a significant effect on the enhancement of the odor response in MBONa1/a2.

### Behavioral role of MBONa1/a2

Mishra et al., (2013) have shown that learnability in *Drosophila larvae* is dependent on the odor intensity at training, i.e. when the concentration at training matches the concentration of testing in an associative learning paradigm, larvae perform more efficient learning compared to lower or higher concentrations of the conditioning odor during testing. Therefore, the learning circuits in the MBs, must have mechanisms to adjust the intensity of odor signal for optimal learning.

The neural basis of associative learning resides at the output synapses between KCs on MBONs that innervate the lobes in a compartment specific manner (Owald et al., 2015). Therefore, these are synaptic sites where learning related plasticity could be induced. In our CATMAID analysis we observed that MBONa1/a2 are presynaptic to many MBONs, as well as modulatory MBINs (Supplementary Fig. 3), suggesting that they can be involved in the regulation of MB output synapses. Could MBONa1/a2 signals regulate KC-MBON synapses to achieve homeostatic control? Homeostatic matching has been reported in flies at the OSN-PN synapse: here strong odor stimuli would cause depression of PN responses to so that PN responses would be transient, a mechanism that would promote perceptual adaptation, and also discriminability of weak signals (Kazama and Wilson, 2008). In this scenario MBONa1/a2, signaling a stimulus intensity largely over its threshold, could depress synapses in the KC output region around the lobes, and modify plasticity and hence learning.

In odor discrimination, subsets of KCs at the core of the MBs, named “on” and “off” cells, have been proposed to function antagonistically to encode increase or decrease of odor intensity, respectively, in an odor discrimination test (Vrontou et al., 2021). They proposed that MBONs innervating the core region of the MBs lobes would be integrating both on and off signals, and the synapses between these both types of KCs and MBONs represent the plasticity site involved in learning; However, on off KCs have yet to be identified. In the larva, there is only one type of gamma KC, and although we can not rule out such “on” and “off” KCs, MBONa1/a2 could potentially be a high-intensity channel and contribute to mediating intensity coding in reinforcement learning.

The only evidence for a role of MBONa1/a2 in odor discrimination comes from studies in Odd neurons by Slater et al., (2015). They found that silencing Odd neurons impairs chemotaxis, while exciting them enhances chemotaxis, and concluded that Odd neurons increase behavioral sensitivity to odor concentrations. Larvae in which Odd neurons were activated could sense 4-fold differences in odor concentration (although at a very high concentration of acid vinegar at 1:4, two to three orders of magnitude higher than the dilutions that we used for learning). Also, when the output of MBONa1/a2 is blocked by expressing *UAS-shi*^*ts*^ in a simplified reward association task, there was an impairment in preference index, the authors have not commented on this, but it might support a role for MBONs in regulating the formation of odor associations. (Saumweber at al., 2018, Supplementary Fig. 5).

### Perspectives

Behavior depends on the precise recognition of sensory cues, however, signals in the environment range not only in its quality but also across different intensity levels. Then, how can an odor object become associated with a given context, even across different intensities?

Here, we found a pair of neurons, MBONa1/a2, that may sense intensity odor signals indirectly via KCs, and that could potentially transmit these signals to output neuropils surrounding the output region of the MBs. Moreover, the prediction is that MBONa1/a2 receive the same regulation as KCs, by receiving inhibitory input via APL, and neuromodulation from octopaminergic sVUM1 neurons. Therefore, MBONa1/a2 would carry not only odor intensity input, but this would be subject to inhibition and modulation, representing a dynamic read-out of the activity state of the calyx upon stimulation of its inputs. Furthermore, since MBONa1/a2 respond only to high concentrations of odor, it could potentially be channeling an odor concentration-dependent pathway that would be integrated with the quality channel of KCs, at the level of the MBONs around the MB lobes. MBONa1/a2 form synapses with many MBONs (Supplementary Fig. 3), and elucidation of downstream partners has the potential to unravel the neural mechanism of the regulation of learning at the MB output synapses. The calyx of the MBs shows remarkable similarity in function and network organization to the piriform cortex of mammals. It is interesting to note that in the mammalian piriform cortex, different separate subsets of piriform cortex neurons have been proposed to carry concentration-invariant odor quality information and concentration-dependent intensity information (Roland et al., 2017). Therefore, the *Drosophila* larval calyx, with a small number of neurons, access to a connectome, and straightforward genetic tools, has the potential to unravel novel universal mechanisms of the regulation of learning.

## Acknowledgements

We thank M Morgan for help in building the optogenetic behavioural apparatus, and Alex McLachlan for technical support. We thank Scott Waddell, Tzumin Lee, Andrew Lin, Kristin Scott and especially the Bloomington *Drosophila* Stock Center for fly stocks, and the Developmental Studies Hybridoma Bank for antibodies. YM and BS were supported by summer research bursaries from Magdalene College of the University of Cambridge. The work was partly supported by BBSRC grant BB/N007948/1 to LMM-N and CJO’K. We thank the Department of Genetics, University of Cambridge, for infrastructural support for LMM-N.

## Author Contributions

LMM-N: Conceived the work. LMM-N, CJO’K, TS, AM: Experimental design. LMM-N, AM, TS, YM, BS: Performed experiments. LMM-N, IM, AM, TS, CJO’K: Data analysis. LMM-N: Paper drafting and writing. All authors: Editing and Reviewing.

## Contribution to the field statement

Learning about the context of a given sensory stimulus with experience, allows an animal to be attracted to food or to escape from danger. The brain therefore has mechanisms to form neural representations of sensory stimuli that are highly selective, to be used to form and retrieve memories. However, the intensity range of sensory signals can be large, in olfaction intensities can vary in the order of 50,000, and it is not clear how odor intensity is integrated to form a selective representation required for memory.

Here we report a pair of neurons in brains of *Drosophila* larva, that receive inputs in the mushroom bodies (MBs), the memory center of the insect brain, that are activated only by high intensity of an odor. We propose that they represent an intensity channel that can be activated in parallel to the “odor quality” channel of MB neurons, and could potentially modify MB downstream memory circuits. This segregation into quality and intensity channels, is also found in the mammalian olfactory cortex, highlighting the potential of *Drosophila* as a model organism, with advanced genetic tools, to unravel universal mechanisms in memory.

