## Supplemental Tables and Figures for "Mushroom body output neurons MBONa1/a2 define an odor intensity channel that regulates behavioral odor discrimination learning in larval *Drosophila*"

**Supplementary Material**

#### Supplementary Tables

**TABLE 1. Presynaptic and postsynaptic sites of MBON-a1/ MBON-a2 in the calyx.** The 3D CATMAID reconstructions of MBON-a1-R, MBON-a2-R, MBON-a1-L and MBON-a2-L from a single 6-hour larva were used to make these measurements.

| Neurons | Number of presynaptic sites in the calyx | Number of postsynaptic sites in the calyx | Total number of synapses in the calyx |
|---|---|---|---|
| MBON-a1-R | 4 | 305 | 309 |
| MBON-a2-R | 2 | 315 | 317 |
| MBON-a1-L | 0 | 222 | 222 |
| MBON-a2-L | 0 | 376 | 376 |

**TABLE 2. Presynaptic and postsynaptic sites of MBON-a1/ MBON-a2 output regions around the MB medial lobe.** 3D CATMAID reconstructions of MBON-a1-R, MBON-a2-R, MBON-a1-L and MBON-a2-L were used to count the number of presynaptic and postsynaptic sites in the ipsilateral and contralateral axonal branches in the output regions. Notice that the number of presynaptic sites is similar to the number of postsynaptic sites.

| Neuron | Number of Presynaptic sites ipsilaterally | Number of Postsynaptic sites ipsilaterally | Number of Presynaptic sites contralaterally | Number of Postsynaptic sites contralaterally |
|---|---|---|---|---|
| MBON-a1-R | 35 | 20 | 38 | 57 |
| MBON-a2-R | 33 | 36 | 38 | 63 |
| MBON-a1-L | 21 | 18 | 39 | 25 |
| MBON-a2-L | 46 | 65 | 35 | 49 |

**TABLE 3. Reciprocal connections among MBON-a1/ MBON-a2 in the output regions.** These were counted using the number of synapses listed for each of the presynaptic partners of each neuron on CATMAID. For example, MBON-a2-R receives input from MBON-a1-R through three synapses, and MBON-a1-L receives input from MBON-a1-R through eight synapses.

| | | Presynaptic Neuron | | | |
|---|---|---|---|---|---|
| | | MBON-a1-R | MBON-a2-R | MBON-a1-L | MBON-a2-L |
| Postsynaptic Neuron | MBON-a1-R | - | 3 | 8 | 1 |
| | MBON-a2-R | 2 | - | 3 | 6 |
| | MBON-a1-L | 4 | 2 | - | 2 |
| | MBON-a2-L | 6 | 5 | 2 | - |

### Supplementary Figure 1

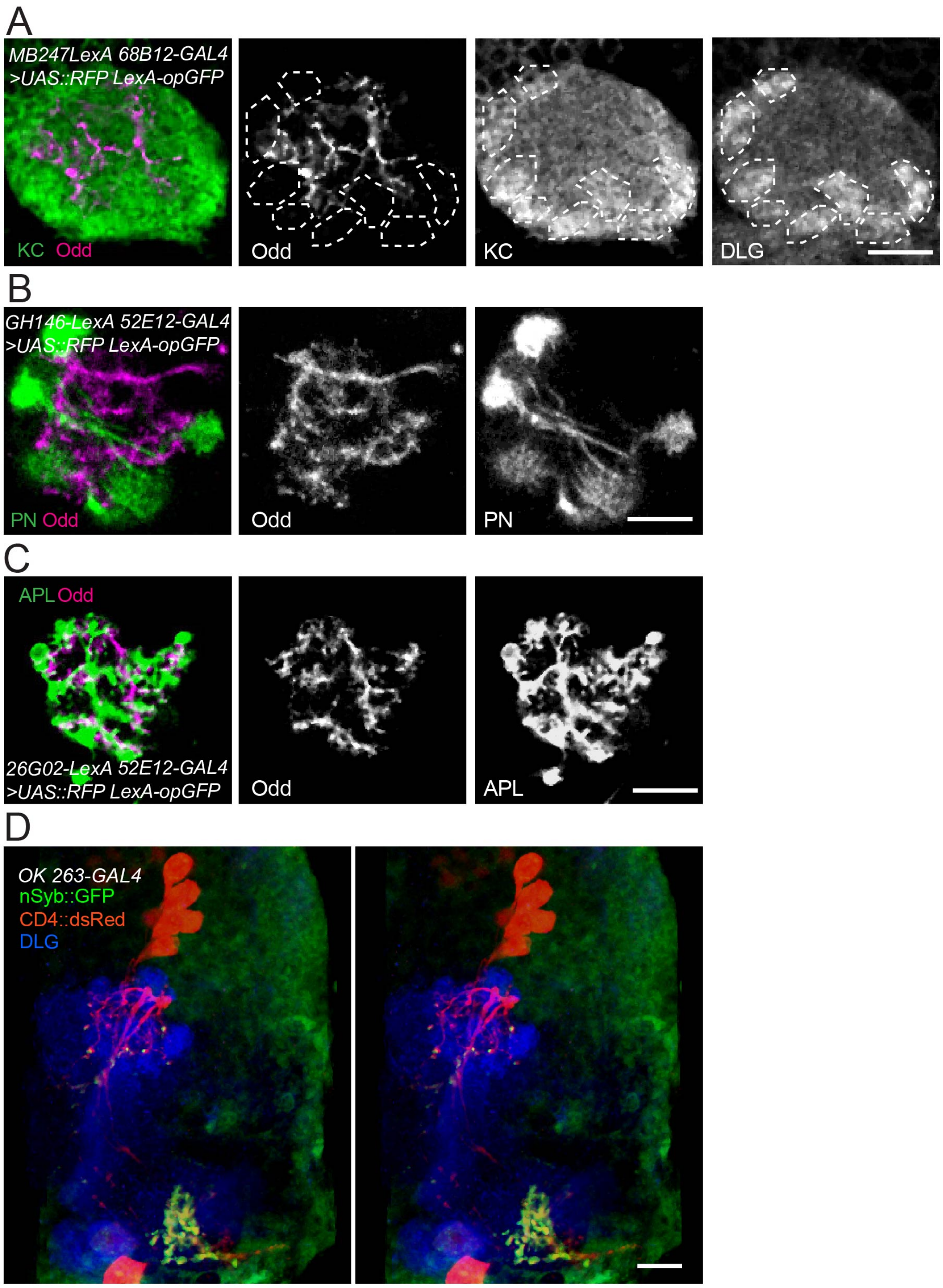


**Supplementary Figure 1. Pattern of calyx innervation by double reporter lines. (A)** Pattern of MBONa1/a2 and KC innervation in the calyx. A line expressing *68B12-GAL4* in a single MBONa1/a2 neuron and *MB247-LexA* in KCs was crossed to reporter line *UAS-mCD8::RFP; LexAop-mCD8::GFP*. Reporter expression was detected by anti-GFP, anti-DsRed and calyx by anti-DLG. Glomeruli are shown by dotted lines. **(B)** MBONa1/a2 expressing *52E12-GAL4* and PNs expressing *GH146-LexA*, showing native fluorescence from the same double reporter. **(C)** APL neuron expressing *26G02-LexA* and MBONa1/a2 expressing *52E12-GAL4,* showing native fluorescence from the double reporter. **(D)** Stereo image of *OK263-GAL4* line labeling MBONa1/a2 with CD4::DsRed and nSyb::GFP, with the MB outline shown by anti-DLG. Panels A-C are single confocal sections of right brain images, anterior to bottom. Panel D is a frontal view of the right brain. Scale bars are 10 µm.

### Supplementary Figure 2

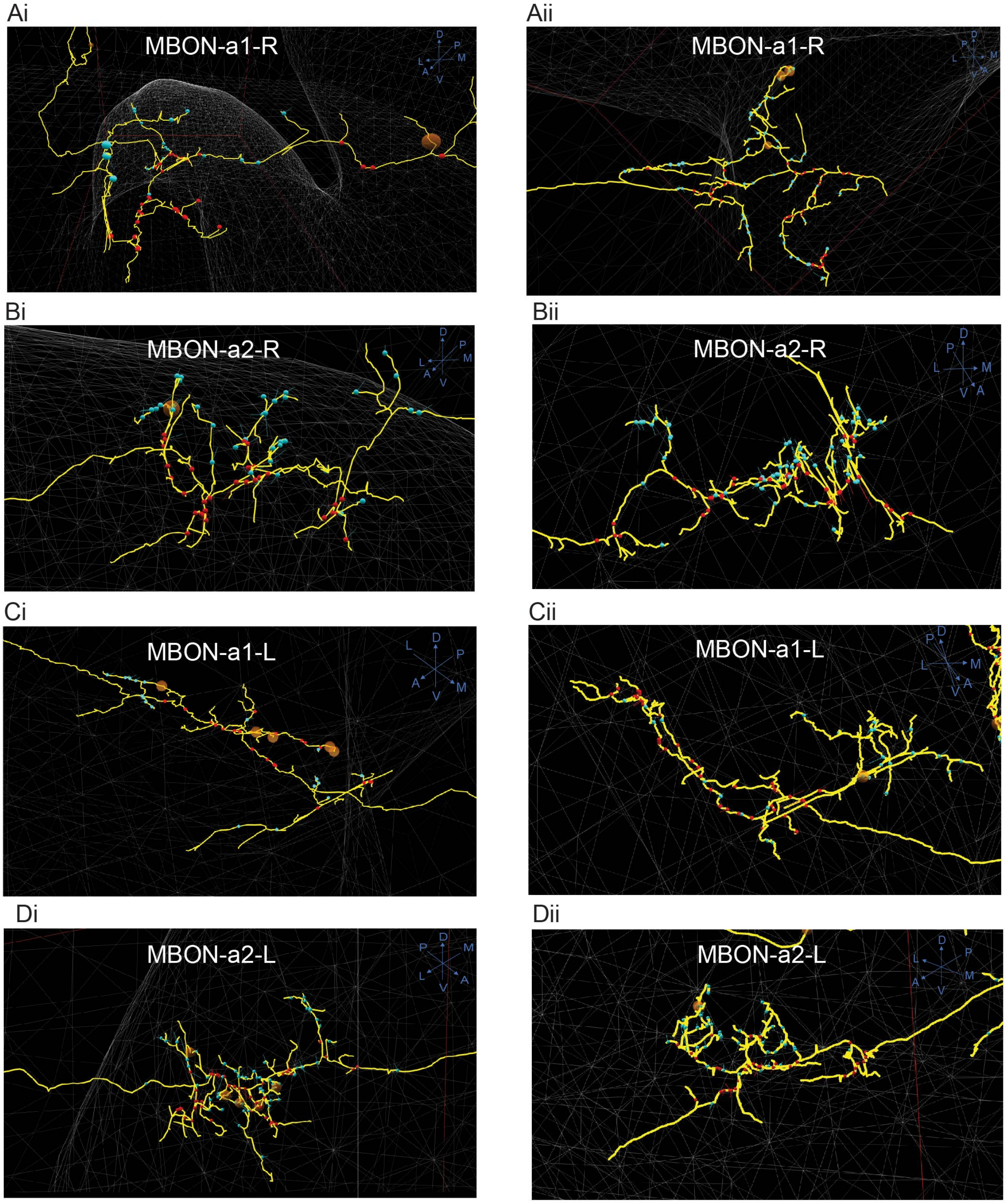


**Supplementary Figure 2. Presynaptic and postsynaptic sites at the ipsilateral and contralateral MB medial lobes visualized using CATMAID.**
Output regions of the four MBON-a1/ MBON-a2 neurons analyzed for synaptic counts. In all panels, small cyan circles are MBON-a1-R and MBON-a2-R postsynaptic sites, and small red circles are presynaptic sites. The larger brown and red circles are unfinished tracing sites. Axes are shown at the top right corner of each panel. Panels show processes of **(A)** MBON-a1-R, **(B)** MBON-a2-R, **(C)** MBON-a1-L and **(D)** MBON-a2-L, all on ipsilateral (i) and contralateral (ii) sides.

### Supplementary Figure 3

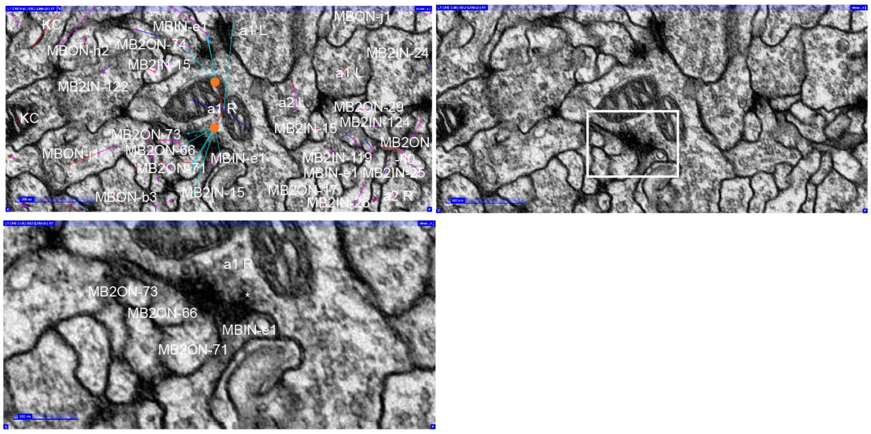


**Supplementary Figure 3**. **Synapses of MBONa1 onto MBONs and MBINs in the output region.**

EM sections of first-instar larva MBONa1/a2. Panels show MBON-a1-L presynaptic to MBON neurons labeled as MB2ONs, and MBINs. The left panel shows an EM section with CATMAID annotations; a connector (orange) is placed on the presynaptic neuron and the cyan arrows indicate postsynaptic partners. The left panel shows the same EM section without the CATMAID annotations, an enlargement of the region within the white square is shown on the bottom panel. Downstream neurons have multiple names which have been omitted in the figure.

MBON-a1-R: https://l1em.catmaid.virtualflybrain.org/?pid=1&zp=39750&yp=36333.76014490835&xp=68737.75884963221&tool=tracingtool&sid0=1&s0=-0.8000000000000008

### Supplementary Figure 4

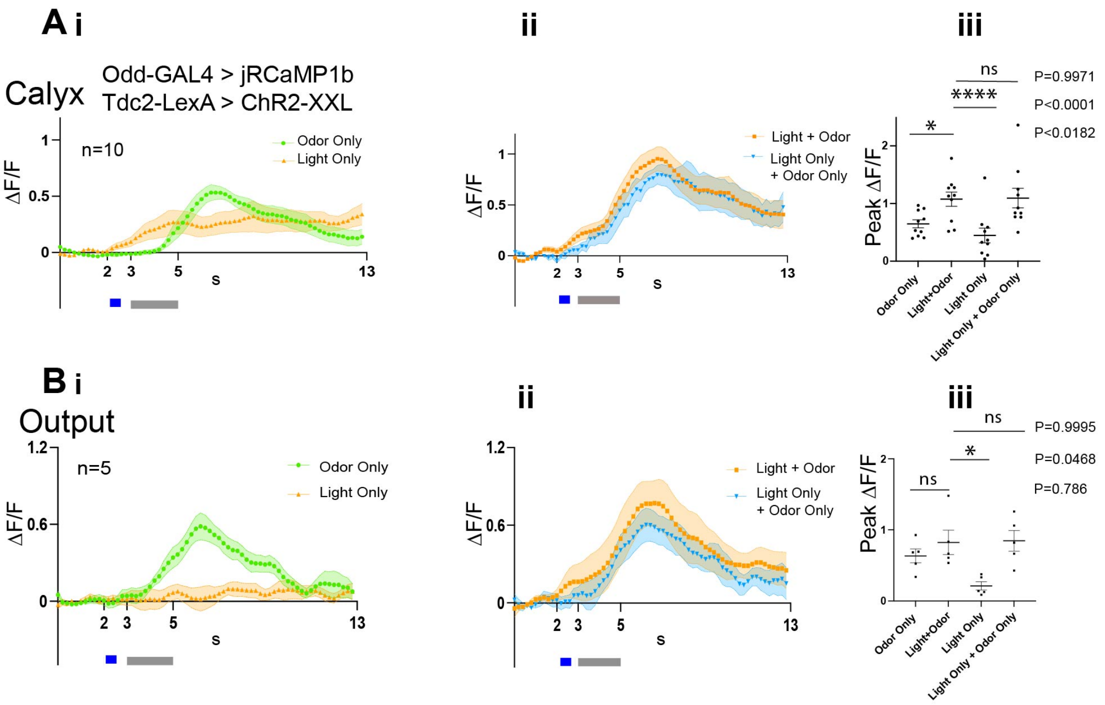


**Supplementary Figure 4. Light contribution to MBONa1/a2 activity in brains expressing ChR2-XXL in *Tdc2-LexA*-expressing neurons.**

To investigate the contribution of the light response to the odor-evoked response in MBONa1/a2 in the larvae used for combined optogenetics and imaging in Fig. 6B and Fig. 6F, we compared the response of MBONa1/a2 to "Light only" and "Odor only" in a set of preparations in which the sequence of (i) Odor only, (ii) Light+Odor was followed by (iii) light only.

**(Ai)** Time courses of MBONa1/a2 calyx ΔF/F from larvae of the same genotype as Fig 6 B and 6F, in response to odor-only or light-only. **(Aii)** MBONa1/a2 responses to odor following activation of ChR2-XXL, and a curve showing the hypothetical ΔF/F values from the sum of odor-only and light-only responses shown in **(Ai). (Aiii)** Comparisons of peak ΔF/F responses for odor only, odor followed by light, light only and the calculated sum of Light only and Odor only responses.

**(B)** Genotype, graphs and analyses are as in **A**, but for the MBONa1/a2 output region.
